# A conserved fertilization complex of Izumo1, Spaca6, and Tmem81 mediates sperm-egg interaction in vertebrates

**DOI:** 10.1101/2023.07.27.550750

**Authors:** Victoria E. Deneke, Andreas Blaha, Yonggang Lu, Jonne M. Draper, Clara S. Phan, Karin Panser, Alexander Schleiffer, Laurine Jacob, Theresa Humer, Karel Stejskal, Gabriela Krssakova, Dominik Handler, Maki Kamoshita, Tyler D.R. Vance, Elisabeth Roitinger, Jeffrey E. Lee, Masahito Ikawa, Andrea Pauli

**Affiliations:** Research Institute of Molecular Pathology (IMP), Vienna BioCenter (VBC), 1030 Vienna, Austria; Vienna BioCenter PhD Program, Doctoral School of the University of Vienna and Medical University of Vienna, Vienna, Austria; Premium Research Institute for Human Metaverse Medicine (PRIMe), Osaka University, Osaka 565-0871, Japan; Department of Experimental Genome Research, Research Institute for Microbial Diseases, Osaka University, Osaka 565-0871, Japan; Institute of Molecular Biotechnology of the Austrian Academy of Sciences (IMBA), Vienna BioCenter (VBC), 1030 Vienna, Austria; Department of Laboratory Medicine and Pathobiology, Temerty Faculty of Medicine, University of Toronto, Toronto, ON, Canada; Laboratory of Reproductive Systems Biology, Institute of Medical Science, The University of Tokyo, Tokyo 108-8639, Japan

## Abstract

Fertilization, the fusion of sperm and egg, is essential for sexual reproduction. While several proteins have been demonstrated to be essential for the binding and fusion of gametes in vertebrates, the molecular mechanisms driving this key process are poorly understood. Here, we performed a protein interaction screen using AlphaFold-Multimer to uncover protein-protein interactions in fertilization. This screen resulted in the prediction of a trimeric complex composed of the essential fertilization factors Izumo1 and Spaca6, and Tmem81, a protein previously not implicated in fertilization. We show that Tmem81 is a conserved, testis-expressed transmembrane protein that is evolutionarily related to Izumo1 and Spaca6 and is essential for male fertility in fish and mice. Consistent with trimer formation *in vivo*, zebrafish *izumo1*^-/-^, *spaca6*^-/-^, and *tmem81*^-/-^ mutants exhibit the same sperm-egg binding defect and show co-depletion of all three proteins in sperm. Moreover, we provide experimental evidence that Izumo1, Spaca6, and Tmem81 interact in zebrafish sperm. Strikingly, the Izumo1-Spaca6 interaction is predicted to form a cleft that serves as a binding site for Bouncer, the only identified egg protein essential for fertilization in zebrafish. Together, these results provide compelling evidence for a conserved sperm factor complex in vertebrates that forms a specific interface for the sperm-egg interaction required for successful fertilization.

## INTRODUCTION

The life of every sexually reproducing organism begins with fertilization, the union of a sperm and an egg. For fertilization to occur, gametes must recognize each other, bind to each other, and fuse. Despite the key importance of this process, our insights into vertebrate sperm-egg binding and fusion lack mechanistic understanding.

Most of our knowledge regarding the molecular basis of vertebrate sperm-egg interaction comes from genetic studies in mice and fish (reviewed in ^1–4^). Over the past 20 years, several proteins have been identified as essential for mammalian gamete interactions. On the sperm, these are the transmembrane proteins IZUMO1^5–7^, SPACA6^8–10^, TMEM95^9,11–13^, FIMP^14^ and DCST1/2^15,16^ as well as the secreted protein SOF1^9^, and on the egg, the tetraspanin proteins CD9/CD81^17–20^ and the GPI-anchored protein JUNO^21^, which directly interacts with IZUMO1^21–24^. Four sperm factors, IZUMO1, SPACA6, DCST1 and DCST2, have known orthologs in fish, and three of them (Spaca6, Dcst1 and Dcst2) have also been shown to be essential for male fertility in fish^16,25^, demonstrating evolutionary and functional conservation of a subset of sperm fertility factors in vertebrates. In contrast, an egg fertility factor conserved across vertebrates has not yet been identified. Mammalian eggs express JUNO^26^, whereas fish eggs express an unrelated Ly6/uPAR protein, Bouncer, which is essential for sperm-egg binding in fish^27,28^. Bouncer is not functionally conserved; its direct mammalian homolog SPACA4 is expressed exclusively in testis and it is required for sperm to penetrate the zona pellucida prior to the contact with the oolemma^29,30^.

While genetic studies have provided a powerful path towards identifying new essential fertilization factors, their molecular functions remain largely unknown. An important exception is the direct interaction between IZUMO1 and JUNO, which has been biochemically and structurally characterized^22–24^. Mammalian cells expressing ectopic IZUMO1 can bind to but not to fuse with JUNO-expressing mouse eggs, which is consistent with a main role for IZUMO1-JUNO in sperm-egg binding^21,31,32^. Although IZUMO1 is similar to SPACA6 and TMEM95 at the structural level^33–35^, recombinant human SPACA6 and TMEM95 do not bind to human JUNO in solution^34,35^. Thus, despite the identification of various essential factors in vertebrates, the precise molecular interactions responsible for sperm-egg binding and fusion remain poorly understood.

The following two questions emerge as the biggest gaps in knowledge in the study of fertilization: Are there essential factors that remain to be identified? And how do these factors contribute collectively to sperm-egg binding and fusion? Here, by employing a protein-protein interaction screen based on AlphaFold structural predictions, we identified Tmem81 as a new essential male fertility factor in fish and mice, and discovered that it interacts with Izumo1 and Spaca6 in mature zebrafish sperm. This trimer provides a molecular connection between the sperm and egg membranes during fertilization by binding to JUNO in mammals and to Bouncer in fish. Overall, our study reveals an intriguing, functionally conserved mechanism of sperm-egg interaction in vertebrates based on the formation of a fertilization complex between egg and sperm.

## RESULTS

### AlphaFold-Multimer predicts the formation of an Izumo1-Spaca6-Tmem81 complex on sperm

To identify putative binding partners of the four known conserved sperm fertility factors in vertebrates, Izumo1, Spaca6, Dcst1 and Dcst2, we performed an AlphaFold-Multimer *in silico* screen^36–38^ against ∼1400 zebrafish testis-expressed secreted/transmembrane proteins (**Fig. 1A**). This approach employs AlphaFold-generated structural predictions of pair-wise protein-protein interactions between each of the four bait proteins and any of the ∼1400 candidate proteins. A high interface predicted template modeling score (ipTM score) reflects high confidence in the relative positioning of the interfaces. Intriguingly, Izumo1-Spaca6 was one of the top-scoring predicted interacting pairs (**Fig. 1B**), which was unexpected given that recombinant human IZUMO1 and SPACA6 ectodomains do not appear to interact *in vitro*^34^. Furthermore, an additional pair amongst the top-scoring predicted pairs, Izumo1-Tmem81, stood out for several reasons. Tmem81 is an uncharacterized, testis-expressed single-pass transmembrane protein that is conserved in vertebrates. Phylogenetic analyses revealed that Tmem81 is related to Izumo1 and Spaca6 (**Fig. 1C, Suppl. Fig. S1A, B**) and shares similarity in the predicted structure (**Fig. 1D, Suppl. Fig. S1D**). The ectodomains of all three proteins contain a hinge region followed by an immunoglobulin (Ig)-like domain, which is followed by a flexible linker and a single, C-terminal transmembrane domain (**Fig. 1D**). The conserved N-terminal four-helix bundle of Izumo1 and Spaca6^34,35^ is absent in Tmem81. Interestingly, apart from Dcst1/2 proteins, which are also present outside of vertebrates, Tmem81 is the most conserved vertebrate fertilization factor identified to date (**Suppl. Fig. S1A**), and its orthologs occur in vertebrates including lampreys, hagfishes and cartilaginous fishes (**Fig. 1C, Suppl. Fig. S1A, B**).

**Figure 1:**
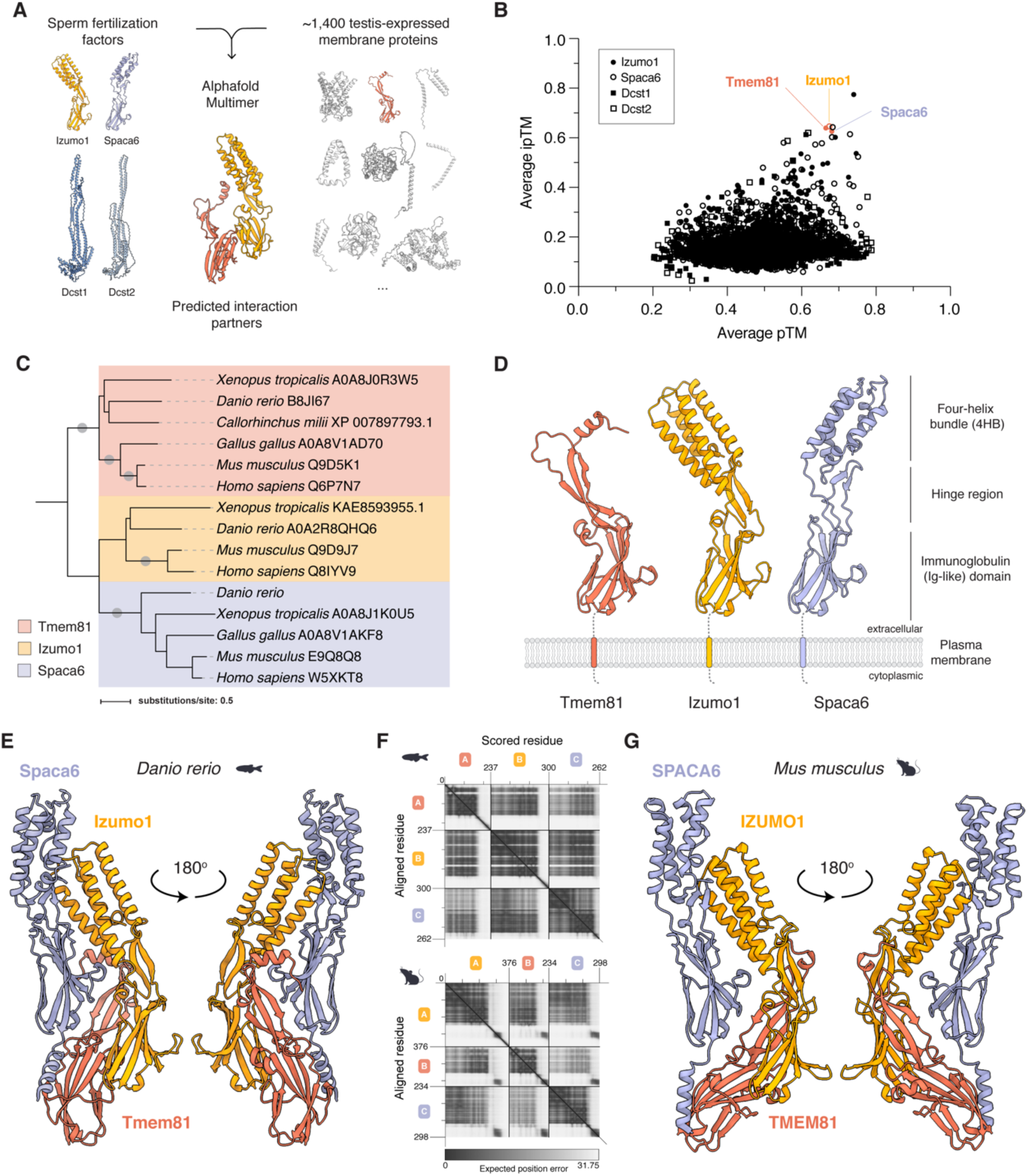
AlphaFold-Multimer predicts the formation of a trimeric complex composed of Izumo1, Spaca6, and Tmem81 on vertebrate sperm. (**A**) Outline of the AlphaFold-Multimer^38,39^ screen to predict putative protein-protein interactions of the four known sperm fertilization factors in zebrafish (baits), Izumo1, Spaca6, Dcst1, and Dcst2, with ∼1400 testis-expressed secreted and membrane proteins (candidates). One resultant predicted interacting pair, Izumo1-Tmem81, is shown as an example. (**B**) Scatterplot of AlphaFold-Multimer predicted ipTM versus pTM scores (ipTM: interface predicted Template Modeling score; pTM: predicted Template Modeling score). The high-scoring pairwise predicted interactions between Izumo1, Spaca6 and the new factor Tmem81 are highlighted in color. (**C**) Phylogenetic tree of the evolutionarily related vertebrate proteins Tmem81 (red), Izumo1 (yellow) and Spaca6 (blue). See **Suppl. Fig. S1B** for an extended tree. (**D**) Tertiary structure predictions of zebrafish Tmem81, Izumo1 and Spaca6. Predictions were generated by AlphaFold2^36^, and protein domains are indicated. (**E**) AlphaFold-Multimer-predicted structural model of the trimeric sperm complex in zebrafish composed of Izumo1 (yellow), Spaca6 (blue) and Tmem81 (red). Structures of the proteins without extracellular linkers and transmembrane domains are shown in two views. (**F**) Predicted Aligned Error (PAE) plots of the top-scoring models of the predicted trimeric complex in zebrafish (top) and mouse (bottom). Each chain represents the full mature protein. Units: amino acid residues; grey scale: expected position error in Angstroms. (**G**) AlphaFold-Multimer-predicted structural model of a trimeric complex in mouse composed of murine IZUMO1 (yellow), SPACA6 (blue), and TMEM81 (red). Structures of the proteins without extracellular linkers and transmembrane domains are shown in two views.

Since Izumo1 was predicted to interact with both Spaca6 and Tmem81 in a pairwise manner (**Fig. 1B**), we assessed whether all three proteins could form a trimer. Indeed, AlphaFold-Multimer^38,39^ predicted with high confidence the formation of a trimer between zebrafish Izumo1, Spaca6, and Tmem81 (**Fig. 1E and F, Suppl. Fig. S1C**). In this trimeric conformation, each of the three proteins, including Spaca6 and Tmem81, are predicted to directly interact with each other such that their Ig-like domains are staggered and align the N-terminal domains facing the extracellular space. Importantly, AlphaFold-Multimer^38,39^ also predicted the formation of a trimer between murine IZUMO1, SPACA6, and TMEM81 (**Fig. 1F and G, Suppl. Fig. S1C**). Together, these results highlighted Tmem81 as a new potential fertilization factor and raised the intriguing hypothesis that a conserved trimeric complex may exist in vertebrate sperm and function during fertilization.

### Tmem81 is essential for male fertility in zebrafish and mice

Tmem81 is an uncharacterized transmembrane protein that shows enriched expression in testis in zebrafish and mammals (**Suppl. Fig. S2A and B**). To study its physiological function in vertebrates, we generated knockout lines in zebrafish and mice using CRISPR/Cas9 (see Materials and Methods; **Suppl. Fig. S2C-F**). In both fish and mice, *Tmem81*^-/-^ animals developed normally and gave rise to adult males and females. However, *Tmem81*^-/-^ male fish and mice were sterile, while females showed normal fertility (**Fig. 2A and B**). Importantly, the male sterility of *Tmem81*^-/-^ mutants was fully rescued in fish and mice by testis-specific transgenic expression of Tmem81-3xFLAG-sfGFP or TMEM81-3xFLAG (Tg), respectively (**Fig. 2A**), revealing that the sterility indeed originated from the absence of Tmem81.

**Figure 2:**
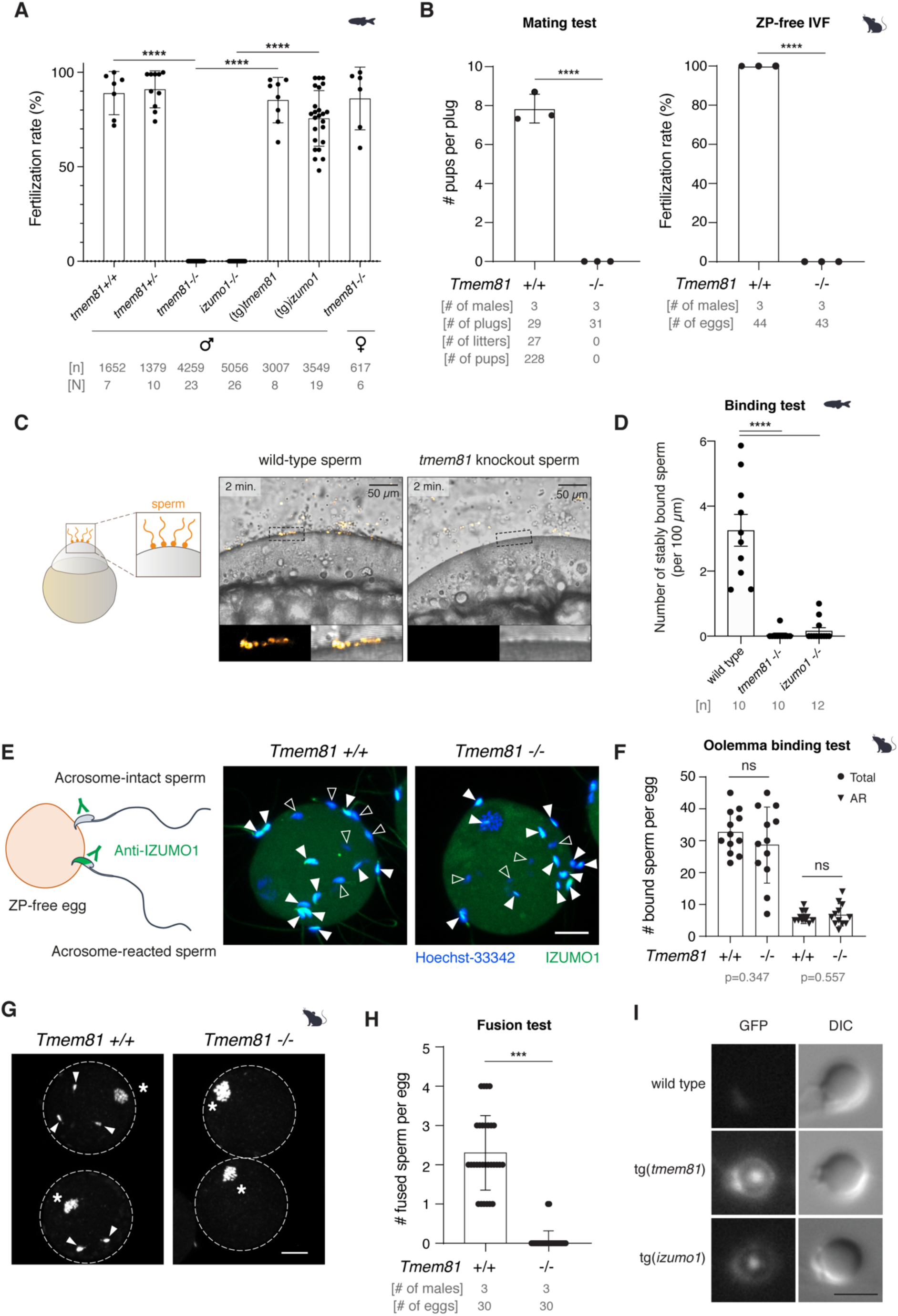
Tmem81 is essential for male fertility in zebrafish and mice. (**A**) Quantification of *in vivo* fertilization rates in zebrafish. Males or females of the indicated genotypes were crossed with wild-type zebrafish of opposite sex. (tg)*tmem81*: *tmem81*^-/-^ fish expressing transgenic Tmem81-3xFLAG-sfGFP; (tg)*izumo1*: *izumo1*^-/-^ fish expressing transgenic Izumo1-3xFLAG-sfGFP. n = total number of embryos; N = number of clutches. Adj. P-value < 0.0001, Kruskal-Wallis test with Dunn’s multiple comparisons test. (**B**) Quantification of *in vivo* (left) and *in vitro* (right) fertilization rates of wild-type and *Tmem81*^-/-^ siblings in mice. For *in vivo* mating tests, wild-type and *Tmem81* knockout males were individually paired with wild-type female mice for eight weeks, during which the copulation plugs and numbers of pups were recorded. For *in vitro* fertilization (IVF) (right), the zona pellucida (ZP) was removed. Numbers of males, plugs, litters, and pups or eggs are indicated. Statistics used an unpaired two-tailed Student’s t-test. (**C**) Sperm-egg binding assay in zebrafish. Left: Experimental setup. Sperm were labeled with MitoTracker (yellow) and incubated with activated and dechorionated wild-type eggs. Right: Representative images of wild-type and *tmem81* knockout (KO) sperm binding to the surface of the egg 2 min after sperm addition. The boxed region is shown at higher magnification below. Scale bar = 50 μm. (**D**) Quantification of wild-type, *tmem81* KO and *izumo1* KO sperm binding to wild-type eggs in zebrafish. Binding of sperm to the egg was quantified by assessing the number of stably bound sperm per 100 μm over a period of 2 min. N = number of independent experiments. Statistics used the Mann-Whitney test. (**E-F**) Sperm–oolemma binding assay in mice. (**E**) Left: Experimental setup. Right: Representative images of wild-type and *Tmem81* KO mouse sperm stained for IZUMO1 (green) and DNA (Hoechst 33342, blue). Filled arrow heads indicate acrosome-reacted sperm, open arrowheads indicate acrosome-intact sperm. Scale bar = 20 μm. (**F**) Quantification of the total number of sperm and the number of the acrosome-reacted (ARed) sperm bound to the oolemma. Ns, not significant. (**G-H**) Sperm–oolemma fusion assay in mice. (**G**) Representative images of the sperm-oolemma fusion assay in mouse. Eggs were pre-loaded with Hoechst 33342 (white). Fused sperm (arrow heads) are highlighted. Asterisks indicate the egg MII chromosomes. Scale bar = 20 μm. (**H**) Quantification of sperm-egg fusion. Statistics used an unpaired two-tailed Student’s t-test. ****: p-value < 0.0001, ***: p-value < 0.001. Means ± SD are indicated. (**I**) Localization of transgenically expressed Tmem81-3xFLAG-sfGFP and Izumo1-3xFLAG-sfGFP in live mature unactivated zebrafish sperm. Scale bar = 3 µm.

Further in-depth analysis of the mutant phenotypes in zebrafish and mice revealed that *Tmem81* knockout (KO) sperm were defective in sperm-egg interaction. In zebrafish *tmem81*^-/-^ mutants, sperm motility was unperturbed and sperm could approach the micropyle (**Suppl. Fig. S2J and S2K**), but failed to stably adhere to dechorionated zebrafish eggs (**Fig. 2C and D**). In mice, *Tmem81*^-/-^ males had normal testis appearance and weight, normal sperm morphology and motility (**Suppl. Fig. S3A-F**), but sperm lacking TMEM81 were unable to fertilize cumulus-intact, cumulus-free, and zona pellucida (ZP)-free eggs *in vitro* (**Fig. 2B, Suppl. Fig. S3G and H**). By monitoring the state of the acrosome through immuno-detection of IZUMO1, we found that sperm lacking TMEM81 could undergo the acrosome reaction and bind the oolemma (**Fig. 2E and F**). However, *Tmem81* KO sperm were defective in fusing with the oolemma of wild-type eggs (**Fig. 2G and H**). Overall, these results identified Tmem81 as a new male fertilization factor essential for sperm-egg interaction in vertebrates.

### The trimer members Izumo1, Spaca6 and Tmem81 interact in mature zebrafish sperm

In line with the predicted trimer formation in mature sperm, we noticed that zebrafish *tmem81*^-/-^ mutants resembled not only the phenotype of the previously characterized *spaca6*^-/-^ mutants in all aspects^25^, but also the defects observed in a newly generated zebrafish line lacking Izumo1 (*izumo1*^-/-^ mutants) (see Materials and Methods and **Suppl. Fig. S2G-I**): In all three mutants, males were sterile and KO sperm failed to bind to the egg (**Fig. 2A, C and D**). Moreover, Tmem81-3xFLAG-sfGFP and Izumo1-3xFLAG-sfGFP showed a similar localization pattern on the head of live mature sperm (**Fig. 2I**). This similar localization pattern on sperm and the same phenotype upon loss of each individual predicted trimer member in zebrafish are consistent with the hypothesis that Izumo1, Spaca6 and Tmem81 may indeed form a trimer on sperm.

If true, stability of each trimer member may depend on the presence of the other two, given that proteins can be destabilized without their binding partners. To test this possibility, we assessed protein abundance in either wild-type or mutant zebrafish sperm lacking one of the three complex members. Shot-gun mass spectrometry (MS) revealed that the vast majority of proteins were unchanged in each mutant compared to wild-type sperm (**Suppl. Table 1**). To confidently detect the three lowly abundant sperm factors, we employed targeted MS via parallel reaction monitoring (PRM)^40^ as a more sensitive method. Although Izumo1, Spaca6, and Tmem81 were confidently and reproducibly detected in wild-type sperm samples, they were either undetectable or substantially reduced (< 10% of wild-type samples) in each of the single mutants (**Fig. 3A**). These results demonstrate that zebrafish *tmem81*^-/-^, *spaca6*^-/-^, and *izumo1*^-/-^ mutants have a common molecular phenotype, consistent with their shared functional phenotypes during fertilization.

**Figure 3:**
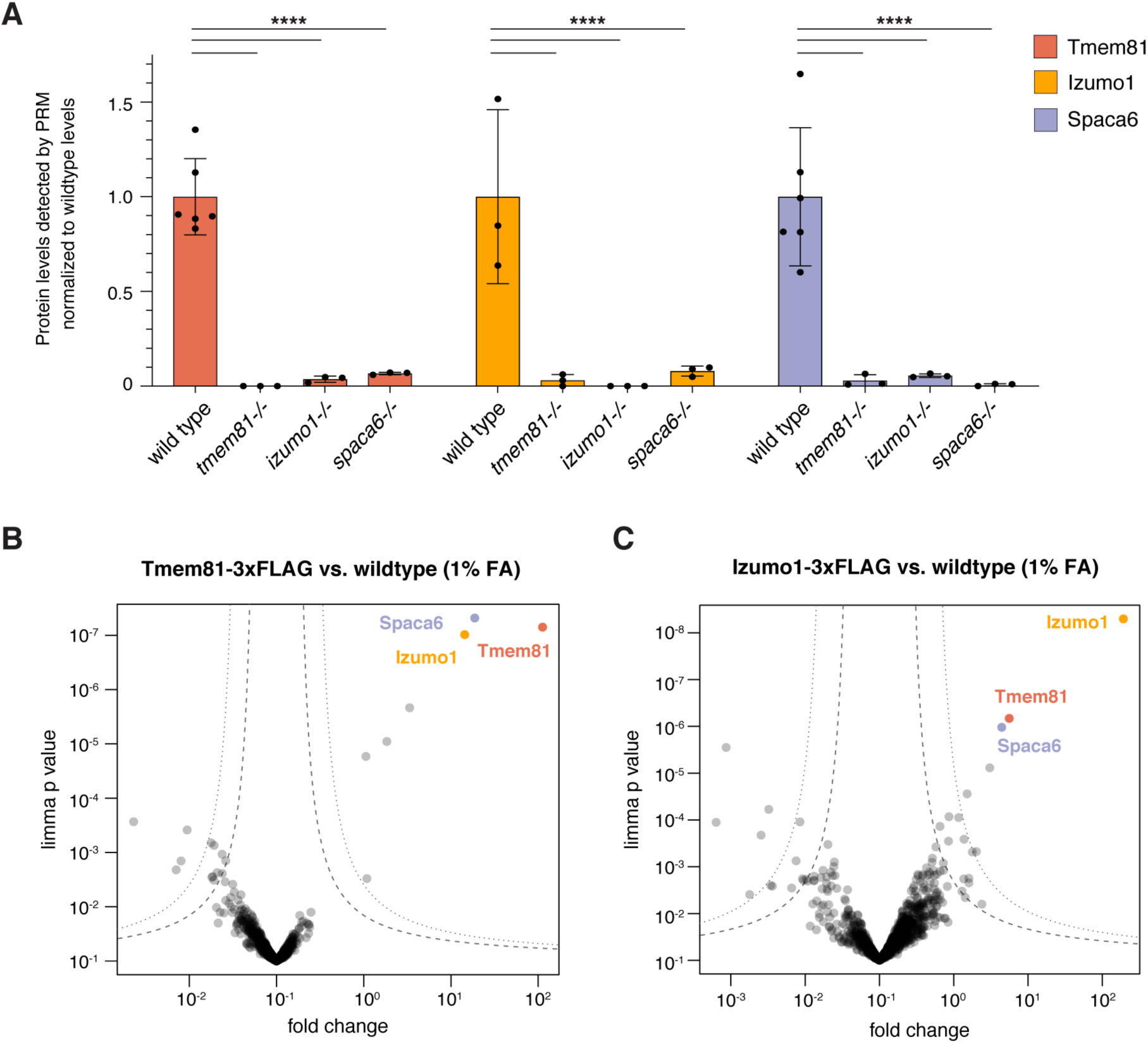
Izumo1, Spaca6, and Tmem81 interact in zebrafish sperm. (**A**) Parallel Reaction Monitoring (PRM)^40^ mass spectrometry quantification of detected protein levels of Tmem81 (red), Izumo1 (yellow) and Spaca6 (blue) in wild-type and mutant zebrafish sperm. Data was normalized by the loaded protein amount determined by 6 constant proteins and is shown relative to the mean wild-type level of each trimer member. ****: p-value < 0.0001. Statistics used the Man-Whitney test. Means ± SD are indicated. (**B**) Volcano plot of differentially enriched proteins in FLAG co-IPs from Tmem81-3xFLAG-sfGFP-expressing sperm versus wild-type sperm. Sperm was lysed after crosslinking with 1% FA (formaldehyde). FLAG co-IPs were performed in triplicates and analyzed by shot-gun MS. Trimeric complex members are highlighted and are the top enriched proteins. (**C**) Co-IPs as in (B) but using Izumo1-3xFLAG-sfGFP-expressing sperm.

The observed loss of all three proteins in each individual zebrafish mutant was in agreement with trimer formation. To investigate the physical interaction between trimer members *in vivo*, we crosslinked sperm before lysis to stabilize protein interactions and performed co-immunoprecipitation followed by mass spectrometry (coIP-MS, see Methods). Notably, Izumo1, Spaca6 and Tmem81 were all enriched by anti-FLAG coIP-MS from Tmem81-3xFLAG-sfGFP versus wild-type sperm (**Fig. 3B**). Performing the reciprocal co-IP with sperm expressing Izumo1-3xFLAG-sfGFP also significantly enriched all complex members, supporting the finding of specific protein-protein interactions between them (**Fig. 3C**).

To address the possibility that we may detect non-specific co-enrichment of membrane proteins, cross-linked co-IP experiments were repeated using sperm expressing the transmembrane domain of human B7 integrin fused to 3xFLAG and sfGFP (TM-3xFLAG-sfGFP)^41^ as an additional negative control. Furthermore, to rule out the possibility of crosslinking artifacts, co-IPs were repeated in the absence of cross-linker. In both control scenarios, all three trimer members were still significantly enriched in the Tmem81-3xFLAG and Izumo-3xFLAG co-IPs (**Suppl. Fig. S4A and B**), validating the specificity of the detected co-enrichment. Taken together, these results support the presence of the predicted trimer of Izumo1, Spaca6, and Tmem81 in zebrafish sperm.

### The Izumo1-Spaca6-Tmem81 sperm complex is predicted to bind Bouncer/JUNO on the egg

A crucial question following the confirmation of the interaction of the trimer members *in vivo* was whether trimer formation could have a direct function in fertilization. A compelling idea was that the Izumo1-Spaca6-Tmem81 trimer could serve as a sperm receptor for proteins exposed on the egg surface, possibly through a composite interaction surface. In zebrafish, Bouncer stood out as the prime candidate because it is on the egg surface, essential for sperm-egg interaction, and restricts fertilization to compatible gametes^27,28,42^. Importantly, the latter directly implies the existence of a Bouncer receptor on sperm.

To determine whether zebrafish Bouncer might indeed interact with the trimer on zebrafish sperm, we used AlphaFold-Multimer^38,39^ to predict complex formation between all four proteins, Izumo1, Spaca6, Tmem81, and Bouncer. Strikingly, Bouncer was predicted to bind to the trimeric complex with high confidence, resulting in the formation of a tetrameric fertilization complex that connects the sperm and egg membranes (**Fig. 4A and B, Suppl. Fig. S5A**). In this model, the cleft between the four-helix-bundles (4HBs) of Izumo1 and Spaca6 at the tip of the trimer forms the binding site for Bouncer, which is predicted to interact through interfaces with the 4HBs of both proteins (**Fig. 4A**). In line with the tetrameric complex bridging two opposing membranes, Bouncer’s GPI membrane anchor is oriented opposite to the transmembrane domains of the sperm proteins (**Fig. 4B**). These structural predictions suggest a key function for the zebrafish sperm trimeric complex in serving as the receptor for Bouncer.

**Figure 4:**
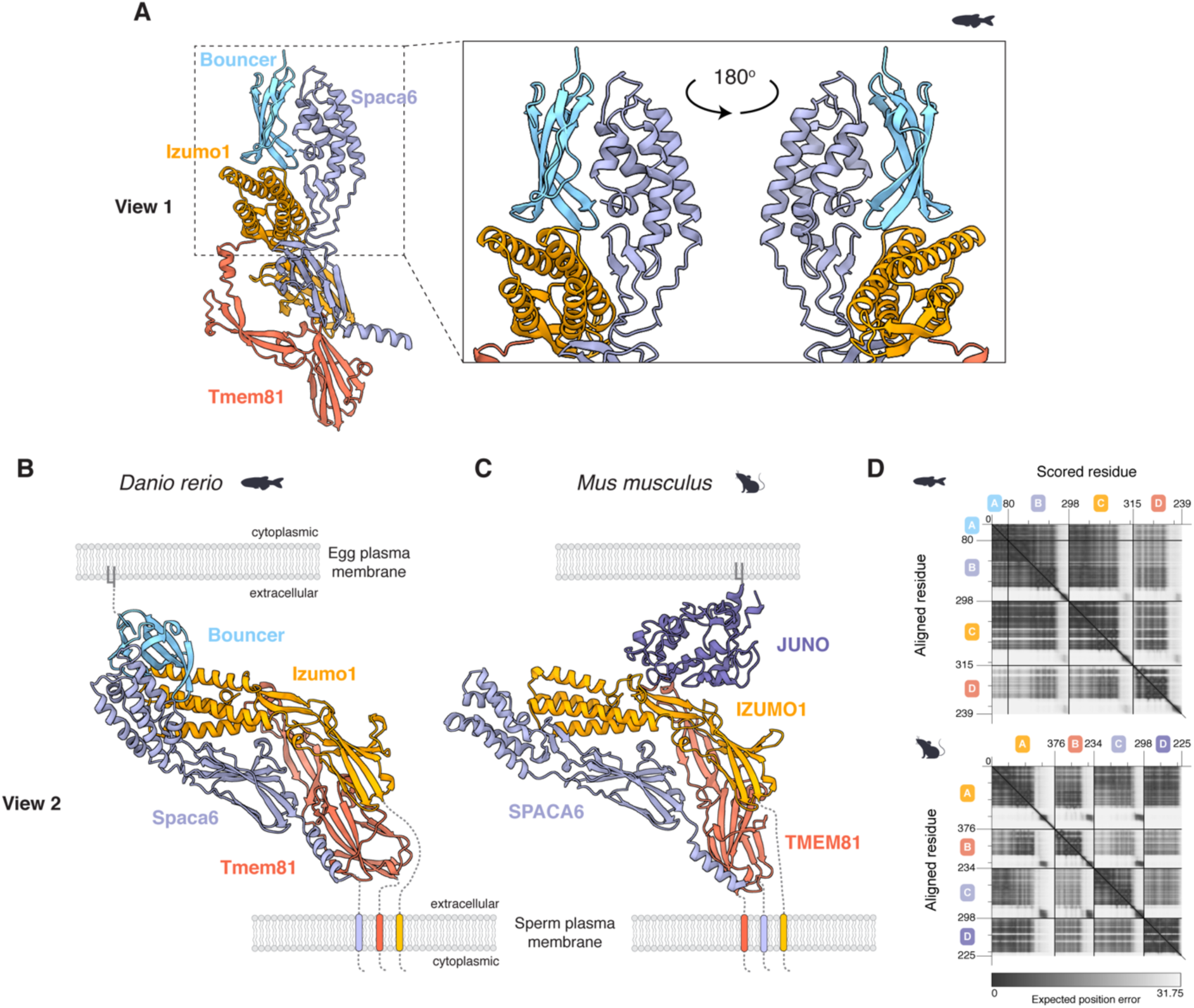
The trimeric sperm complex of Izumo1, Spaca6, and Tmem81 is predicted to bind to Bouncer/JUNO on the egg. (**A**) AlphaFold-Multimer predicted structural model of a tetrameric complex formed by binding of the egg protein Bouncer (light blue) to the trimeric complex on sperm composed of Izumo1 (yellow), Spaca6 (blue) and Tmem81 (red). A higher magnification view of the boxed area is shown on the right in two views, highlighting the binding site of Bouncer in the cleft between Izumo1 and Spaca6 that is formed at the tip of the trimer. (**B**) Rotated view of the predicted tetrameric complex shown in (A), bridging sperm and egg membranes in zebrafish. (**C**) AlphaFold-Multimer predicted structural model of a tetrameric complex bridging sperm and egg membranes in mice. In mice, JUNO (dark purple) binds to IZUMO1^22–24^, and this interaction is predicted to be compatible with trimer formation on murine sperm. (**D**) Predicted Aligned Error (PAE) plots of the top-scoring models of the predicted trimeric complex with Bouncer in zebrafish (top) and with JUNO in mouse (bottom). Each chain represents the full mature protein. Units: amino acid residues; grey scale: expected position error in Angstroms. In (A)-(C), structures are shown as cartoons.

In contrast to Bouncer, its mammalian homolog, SPACA4, is expressed in testis and not in oocytes^29,30,43^, and is not predicted to interact with the mammalian IZUMO1-SPACA6-TMEM81 trimer by AlphaFold-Multimer despite sharing Bouncer’s Ly6/uPAR protein fold^27^. However, in mammals, IZUMO1 is known to directly bind to GPI-anchored JUNO on the egg^21–24^. AlphaFold-Multimer, which is trained on solved structures of protein-protein interactions including the published structures of the human IZUMO1-JUNO complex^22–24^, predicted IZUMO1-JUNO binding to be compatible with trimer formation (**Fig. 4C, Suppl. Fig. S5B**). Thus, the conserved function of the trimer on vertebrate sperm may be to connect sperm and egg membranes by interacting with a ligand on the egg that has diverged during vertebrate evolution.

## DISCUSSION

Most of the current knowledge in the fertilization field is based on genetic studies, which have identified all essential fertilization factors so far. However, insights into the molecular mechanisms of these factors have been lagging behind. While prior studies suggested that fertilization factors may form protein complexes^16,44,45^, no direct evidence apart from the interaction between mammalian IZUMO1 and JUNO exists to date^22–24^. Through structural predictions supported by direct experimental evidence *in vivo*, our study reveals the formation of a conserved trimeric complex on sperm in zebrafish, which is also predicted to form in mammals. This complex is composed of the two known vertebrate male fertility factors Izumo1 and Spaca6 and the newly identified related factor Tmem81 that we demonstrate to be essential for male fertility in zebrafish and mice. Remarkably, trimeric complex formation is predicted to generate the binding site for zebrafish Bouncer, thereby forming a tetrameric protein complex connecting the membranes of sperm and egg and bringing them in close proximity. In mammals, JUNO’s binding site on the hinge of IZUMO1 is accessible in the predicted mammalian trimer. Hence, Bouncer may have a functionally equivalent role to JUNO in mediating adhesion, and potentially fusion, of compatible sperm and egg by binding the conserved sperm trimeric complex.

Interestingly, JUNO and Bouncer are both GPI-anchored proteins that arose by gene duplications and are expressed in the egg of mammals or fish, respectively, but are evolutionarily unrelated and structurally distinct proteins^21,27^. JUNO originated from a mammalian-specific gene duplication of the Folate-receptor 4 gene^26^, while Bouncer belongs to the large class of Ly6/uPAR proteins, which have undergone multiple gene duplications in vertebrates and share a characteristic three-finger-fold^27,46^. Gene duplications are generally thought to be a primary mechanism through which new genes and new biological functions arise^47,48^, driving phenotypic diversity across the tree of life. In light of this, the proposed formation of a tetrameric complex composed of conserved factors on the sperm and a divergent factor on the egg poses an interesting conundrum due to the presumed analogous function fulfilled by the egg factor. A possible explanation for this phenomenon could be that evolution of new ligands on the egg, which can bind to distinct regions of the trimer, could accommodate the high environmental diversities during vertebrate fertilization. Given the distinct binding sites on the trimer, we expect the interacting residues to be under differential selective pressure to co-evolve in the different vertebrate phyla and contribute to species-specific fertilization. This phenomenon has already been described for the interaction between IZUMO1 and JUNO^26^ and has also been observed in the case of Bouncer between zebrafish and medaka, though here the sperm interaction partners were still unknown^27,28^. Overall, our work suggests evolutionary diversification of the interaction partner of the trimer on the egg while all three sperm proteins constituting the trimer are widely conserved in vertebrates. Our identification of the conserved trimer and similar predictions performed by Jovine and colleagues^49^ argue for a striking level of conservation at the level of protein identity and complex formation of a putative core machinery mediating fertilization on sperm. Interestingly, the study by Jovine and colleagues raises the possibility that, in addition to JUNO, CD9 on the mammalian egg may bind the mammalian trimeric complex at the same site as Bouncer in fish^49^. Future work will be required to test these hypotheses and experimentally validate the binding of the proposed egg interacting factors to the trimer.

Our study suggests that the main function of the trimeric complex on sperm is to provide a receptor for ligands on the egg, thereby facilitating the formation of a ‘fertilization synapse’^50^. While knockout of any one factor leads to reduced abundance of the other two in zebrafish sperm, published data suggests that this phenomenon is not universally conserved because at least IZUMO1 protein persists in murine sperm lacking SPACA6^50^. Interestingly, members of the trimeric complex may also have functions outside of the complex. This has been suggested for IZUMO1, which has been proposed to dimerize upon oocyte binding^45^ and has been reported to fuse somatic cells overexpressing solely IZUMO1 and no other trimeric complex member^31^, is currently unclear. Whether and how the contact between the trimer and the egg is involved in fusion remains to be studied.

## ACKNOWLEDGEMENTS

We thank Juraj Ahel for setting up a pipeline for running Alphafold-Multimer on our local cluster; Clemens Plaschka for advice on the analysis of the predicted structures; David Sanders for sharing a plasmid containing the B7 integrin transmembrane domain; Maria Novatchkova for help with generating a list of testis-expressed secreted/transmembrane-containing proteins in zebrafish; all members of the proteomics facility at IMP/IMBA/GMI using the VBCF instrument pool for their continuous support of our work; all members of the bio-optics facility of the Vienna Biocenter Core Facilities (VBCF) for their continuous support; Luca Jovine for sharing his pre-print prior to submission to bioRxiv; Krista Gert, Laura Lorenzo-Orts, Josef Roehsner, Ulrich Hohmann and XinYin for valuable discussions and support; Verena Ruprecht and Yanjun Liu for continuous discussions; the fish facility personnel and aquatics team for taking care of zebrafish; Mirjam Binner, Anna Bandura, Tetiana Mylenko, and Gisela Deneke for their help with genotyping; David Drechsel and his team at the Protein Technologies facility of the Vienna Biocenter Core Facilities (VBCF) for their continuous support; Egon Ogris and Stefan Schuechner as well as their team of the monoclonal antibody facility at the Max Perutz Labs for their support; Enrica Bianchi and Gavin Wright for discussions and support; Angela Andersen from the LifeScience editors for valuable feedback on the manuscript; and the entire Pauli lab for helpful discussions about the project and feedback on the manuscript.

## AUTHOR CONTRIBUTIONS

V.E.D., A.B. and A.P. conceived the study; V.E.D. and A.B. designed, performed, and analyzed experiments; Y.L. and M.I. led the mammalian work with help from M.K.; J.M.D., C.S.P., K.P., L.B. and T.H. contributed to the experimental work and analyses in zebrafish; A.S. conducted phylogenetic analysis and discovered Tmem81; K.S. and G.K. performed the PRM mass spectrometry experiments; E.R. supervised and coordinated the mass-spectrometric analyses; D.H. established the AlphaFold-Multimer batch pipeline on our local cluster; T.D.V. performed and J.E.L. supervised biochemical experiments; A.P. supervised the study; V.E.D., A.B. and A.P. wrote the manuscript with contributions from Y.L. and input from all authors.

## MATERIAL AND METHODS

### AlphaFold-Multimer predictions

To predict protein-protein interactions, Uniprot protein sequences were obtained and listed in FASTA format. A ‘baits’ file was generated with the sequences of the known zebrafish fertilization factors Izumo1 (Uniprot ID: A0A2R8QHQ6), Spaca6 (NCBI: NP_001384707.1), Dcst1 (Uniprot ID: A0A8M6YVA5) and Dcst2 (Uniprot ID: A0A8M6Z1B9) excluding their signal peptide sequences. A ‘candidates’ file was created by filtering all testis-expressed genes (based on RNA-sequencing data from ^27^) for proteins that contain transmembrane and/or signal peptide sequences (∼1400 candidates), and signal peptide sequences were removed for the screen. Alphafold-Multimer^36–38^ was used to predict interactions between the ‘bait’ proteins and ‘candidate’ proteins with a script provided by D. Handler using mmseqs^51^ (git@8c41480) for local MSA creation and colabfold^38^ (git@38e21e3) for structure prediction.

### Analysis of the evolutionary relationship between Izumo1, Tmem81 and Spaca6

Tmem81 (Uniprot ID: B8JI67) was independently discovered in a search for remote homologs of Spaca6 proteins by an iterative PSI-BLAST in the UniProt reference proteomes with a restrictive E-value threshold of 1e-5 with the mature zebrafish Spaca6 excluding the signal peptide (NCBI protein accession: NP_001384707.1:19-317)^52–54^. Spaca6 orthologs were collected in round 1 and 2. In round 3, Izumo1 proteins were included, such as human IZUMO1 (Q8IYV9, E-value 1.71e-35). In the same round with higher E-values but still highly significant, an additional protein family appeared, Tmem81, represented by human TMEM81 (Q6P7N7, E-value 2.46e-15). Similar results were obtained with other members of the Spaca6 family. A maximum likelihood phylogenetic tree of Spaca6, Izumo1 and Tmem81 proteins was calculated with IQ-TREE 2 (v.2.2.0^55^), using an alignment of selected family members with mafft (v7.505, -linsi method^56^), extracting the conserved region covering zebrafish Spaca6 from residues 83 to 307 with Jalview^57^, and restricting the alignment to columns covering more than 20% of the sequences. The phylogenetic tree was inferred with standard model selection using ModelFinder^58^ and ultrafast bootstrap (UFBoot2) support values^59^. The tree was visualized in iTOL (v6^60^). Branches that are supported by an ultrafast bootstrap value >=95% are indicated by a grey dot. Branch lengths represent the inferred number of amino acid substitutions per site, and branch labels are composed of gene name (if available), genus, species, and accession number.

To generate a taxonomic tree of fertility factors conserved across vertebrates, orthologs of all protein families were collected in individual blast searches (no iterations) starting with zebrafish or human representatives in the NCBI nr protein database and UniProt reference proteomes applying significant E-value thresholds (< 1e-4). Proteins with the DC_STAMP domain were determined in a search with the PFAM HMM PF07782^61^. Significant hits of selected species were aligned with mafft (-linsi)^56^ and the conserved region covering residues 48 to 607 of human DCST1 (UniProt: Q5T197) was extracted with Jalview^57^. A maximum likelihood phylogenetic tree was inferred with IQ-TREE for classification of the Dcst1 and Dcst2 subfamilies. The taxonomic tree was generated with the NCBI Taxonomy Common Tree webtool and visualized with iTOL.

### Zebrafish husbandry and ethics statement

Zebrafish (*Danio rerio*) were raised according to standard protocols with a 14/10h light/dark cycle at 28 °C water temperature. The wild type zebrafish were acquired by crossing TL (Tupfel Longfin) and AB zebrafish. All zebrafish experiments were conducted according to Austrian and European guidelines for animal research and approved by the Amt der Wiener Landesregierung, Magistratsabteilung 58—Wasserrecht (zebrafish protocols GZ 342445/2016/12 and MA 58-221180-2021-16).

### Generation of zebrafish mutants for *tmem81* and *izumo1*

Zebrafish knock-out fish for *tmem81* and *izumo1* were generated by CRIPSR/Cas9-mediated mutagenesis according to standard protocols^62^. In brief, 2 guide RNAs (gRNAs) targeting first coding exons of *tmem81* or *izumo1* were generated according to published protocols^62^ by oligo annealing followed by T7 polymerase-driven *in vitro* transcription (gene-specific targeting oligos: tmem81_gRNA: 5’-TACGTGAGACAACCCAGGTGTGG-3, 5’-ACTCAAGAACTCTGTCCTATTGG-3’; izumo1_gRNA: 5’-ACTCGATCTGATCACGGACGCGG-3’, 5’-CATGTACATAACGGATGACCTGG-3’; common tracer oligo AAAAGCACCGACTCGGTGCCACTTTTTCAAGTTGATAACGGACTAGCCTTATTTTAACTTGCTATTTCTAGCTCTAAAAC). Cas9 protein and gRNAs were co-injected into the cell of one-cell stage TLAB embryos. Putative founder fish were outcrossed to TLAB wild-type fish. Founder fish carrying germline mutations were identified by size differences in the gene-specific amplicons in a pool of embryo progeny (primers: tmem81_gt_F: 5’-ATCTACTGACAGCTTGGCTAGAC-3’ and tmem81_gt_R: 5’-TTGTCCAGGCTAAGGGCTTC-3’; izumo1_gt_F: 5’-TTCACTCCAGTCTCTCCCAAAT-3’ and izumo1_gt_R: 5’-TCGGATTTACCTATAACACCGC-3’). Embryos from founder fish were raised to adulthood. Heterozygous males and females of each mutant genotype were incrossed to generate homozygous mutant embryos. Sanger sequencing of homozygous mutant embryos identified the nature of the mutation. The *tmem81* mutation is a 119-nt deletion in exon 2, which results in a frameshift mutation and a premature stop codon after 92 amino acids. The *izumo1* mutation is a 14-nt insertion in exon 1, which results in a frameshift mutation and a premature stop codon after 55 amino acids. Genotyping of mutant fish was performed by PCR, and detection of the mutant product was performed by standard gel electrophoresis using a 4% agarose gel.

### Generation of zebrafish transgenic lines

Zebrafish transgenic lines expressing Tmem81-3xFLAG-sfGFP and Izumo1-3xFLAG-sfGFP under the control of the testis-specific *dcst2* promoter^16^ were generated by Tol2-mediated transgenesis. In brief, *tmem81*^+/-^ or *izumo1*^+/-^ zebrafish embryos were injected at the 1-cell stage with 35 ng/μL *tol2* mRNA, 15 ng/μL of the respective expression construct and 0.083% phenol red solution. The embryos were kept in E3 medium at 28 °C and those with strong somatic expression of the transgenesis markers were considered putative founders and raised to adulthood. These fish were crossed with *tmem81*^+/-^ or *izumo1*^+/-^ zebrafish and the progeny was screened for the transgenesis marker indicating stable germline transmission. The transgenic progeny was raised to adulthood and genotyped to identify transgenic fish in a homozygous mutant (*tmem81*^-/-^ or *izumo1*^-/-^) background.

Zebrafish transgenic lines expressing transmembrane (TM)-3xFLAG-sfGFP under the control of the testis-specific *dcst2* promoter^16^ were generated as described above, but in a wild-type TLAB zebrafish background.

### Fertility assessment of adult zebrafish

Zebrafish *in vivo* fertilization rate measurements were performed by counting fertilization rates of natural matings. The evening prior to mating, the fish assessed for fertility and a TLAB wild-type fish of the opposite sex were separated in breeding cages. The next morning, the fish were allowed to mate. Eggs were collected and kept at 28 °C in E3 medium (5 mM NaCl, 0.17 mM KCl, 0.33 mM CaCl_2_, 0.33 mM MgSO_4_, 10^-5^% Methylene Blue). The rate of fertilization was assessed approximately 3 hours post-laying. By this time, fertilized embryos have developed to ∼1000-cell stage embryos, while unfertilized eggs resemble one-cell stage embryos.

### Zebrafish sperm and egg collection for *in vitro* assays

Zebrafish unactivated gametes were collected as previously described^27,28^. In brief, the evening prior to sperm and egg collection, male and female zebrafish were separated in breeding cages (one male and one female per cage). To collect unactivated sperm, male zebrafish were anesthetized using 0.1% tricaine, and unactivated sperm was collected in a capillary by mouth pipetting under a dissecting microscope. The sperm was stored on ice in Hank’s saline solution (5.4 mM KCl, 0.137 M NaCl, 1 mM MgSO_4_, 4.2 mM NaHCO_3_, 0.25 mM Na_2_HPO_4_, 1.3 mM CaCl_2_). To collect unactivated eggs, female zebrafish were anesthetized using 0.1% tricaine, and eggs were expelled into a dry petridish by gentle pressure on the belly of the female.

### Localization of transgenic GFP-tagged Tmem81 and Izumo1 in zebrafish sperm

Sperm from transgenic Tmem81-3xFLAG-sfGFP, Izumo1-3xFLAG-sfGFP males or wild-type males were collected in Hank’s saline, dropped onto a glass slide and covered with a cover slip. Samples were imaged with an Axio Imager.Z2 microscope (Zeiss) using an oil immersion 100x/1.4 plan-apochromat objective and recorded with a Hamamatsu Orca flash 4 v3 camera followed by processing using Fiji by adjusting image brightness and contrast.

### Sperm egg-binding assays in zebrafish

Sperm-egg binding assays using activated, dechorionated eggs were performed as previously described^16,25^. In brief, unactivated sperm was collected from 2 to 4 fish per genotype and kept in 100 μL Hank’s saline + 0.5 μM MitoTracker DeepRed^FM^ on ice. Un-activated, mature eggs were collected from a wild-type female fish and activated by addition of E3 medium. After 10 min, 1–2 eggs were manually dechorionated using forceps and transferred to a cone-shaped imaging dish with E3 medium. After focusing on the egg plasma membrane, the objective was briefly lifted to add 2–10 μL of stained sperm (∼200,000–250,000 sperm). Imaging was performed with a LSM800 Examiner Z1 upright system (Zeiss) using a 10x/0.3 Achroplan water dipping objective. Images were acquired until sperm were no longer motile (5 min). To analyze sperm-egg binding, stably-bound sperm were counted. Sperm were counted as bound when they remained in the same position for at least 1 min following a 90-s activation and approach time window. Data was plotted as the number of sperm bound per 100 μm of egg membrane for one minute.

### Sperm approach and sperm motility assays in zebrafish

Sperm approach and sperm motility assays were performed as previously described^16,25^.

### Mouse husbandry and ethics statement

B6D2F1/J and ICR mice were purchased from Japan SLC, Inc. (Shizuoka, Japan) and maintained under cycles of 12 h light and 12 h darkness with ad libitum feeding. All animal experiments were approved by the Animal Care and Use Committee at the Research Institute for Microbial Diseases, Osaka University (#Biken-AP-H30-01) and were conducted in compliance with all guidelines and regulations. Mice were euthanized by cervical dislocation following anesthesia.

### Generation of a mouse *Tmem81* knockout and TMEM81-3xFLAG transgenic line

A *Tmem81* knockout mouse line was generated by CRIPSR/Cas9 using two single guide RNAs (sgRNAs) flanking the coding region of *Tmem81*. Wild-type (B6D2F1/J) female mice were superovulated by peritoneal injection of CARD HyperOva (Kyudo Co., Saga, Japan) and human chorionic gonadotropin (hCG; ASKA Pharmaceutical Co. Ltd., Tokyo, Japan), and were individually paired with a wild-type male. Two-pronuclear (2PN) zygotes were collected from the female mice. A ribonucleoprotein complex of CRISPR RNA (crRNA), trans-activating crRNA (tracrRNA), and Cas9 was introduced into the 2PN eggs using a NEPA21 super electroporator (NEPA GENE, Chiba, Japan). The treated zygotes were cultured in the potassium simplex optimization medium (KSOM) to two-cell stage and transplanted into the ampullary segment of the oviducts of 0.5-day pseudo-pregnant ICR females. The founder mice were obtained by natural delivery or Cesarean section and were genotyped by PCR. The mutant allele was subsequently examined by Sanger sequencing.

To produce a transgenic TMEM81-3xFLAG (Tg) expressing mouse line, the open reading frame of *Tmem81* was cloned from mouse testis cDNA by PCR and inserted into a plasmid encoding an Izumo1 promoter and a rabbit beta globin polyadenylation (polyA) signal. The linearized plasmid was microinjected into the pronuclei of zygotes obtained from mating between homozygous knockout females and heterozygous knockout males. The injected zygotes were cultured in the KSOM medium until the two-cell stage and transplanted into the oviductal ampulla of 0.5 d pseudo-pregnant ICR females. Founder animals were obtained by natural delivery or Cesarean section after 19 days of pregnancy. The transgene was identified by PCR using primers targeting the Izumo1 promoter and the polyA signal.

Frozen spermatozoa from *Tmem81*^+/–^ males (B6D2-Tmem81<EM1OSB>), and *Tmem81*^+/–^ Tg males (B6D2-Tg(*Izumo1-Tmem81/3×FLAG*)1Osb) will be available at the Riken BioResource Center (RIKEN BRC; web.brc.riken.jp/en) and the Center for Animal Resources and Development, Kumamoto University (CARD R-BASE; cardb.cc.kumamoto-u.ac.jp/transgenic).

### Mating tests

Upon sexual maturation, *Tmem81*^-/-^ or *Tmem81*^-/-^ Tg male mice were caged individually with three 6-week-old wild-type female mice for eight weeks. During this period, vaginal plugs was examined as an indicator of successful copulation and the number of pups was counted at birth. Three knockout males were tested to meet the requirements for statistical validity. The fecundity of three wild-type males was tested in parallel as positive controls. After the eight-week mating period, the male mice were withdrawn from the cages and the females were kept for another three weeks to allow their final litters being delivered.

### Sperm morphology and motility

Cauda epididymal sperm were extracted from adult male mice under a dissection microscope using micro spring scissors and fine forceps. For analyzing sperm morphology, sperm were suspended in 1 mL PBS by gentle pipetting and were observed under an Olympus BX-53 differential interference contrast microscope equipped with an Olympus DP74 color camera (Olympus, Tokyo, Japan). For motility analysis, sperm were dispersed in 100 μL of Toyota, Yokoyama, and Hosi (TYH) medium drops. Sperm motility was measured by the CEROS II sperm analysis system (Hamilton Thorne Biosciences, Beverly, MA) at 10 min and 2 h of incubation in TYH medium.

### *In vivo* fertilization

Wild-type female mice were superovulated by peritoneal injection of PMSG and hCG and paired with wildtype or *Tmem81*^-/-^ males. Oviducts were isolated from the female mice 6 h after the formation of copulation plugs. COCs extracted from the oviductal ampulla were incubated in the KSOM medium and treated with 330 μg/mL of hyaluronidase (Wako, Osaka, Japan) to remove the cumulus cells. The eggs were then fixed on ice in FHM medium containing 0.25% (v/v) glutaraldehyde (Polysciences, Inc., Warrington, PA) for 15 min, washed in fresh FHM medium for three times, and stained with Hoechst 33342 at room temperature for 15 min to visualize pronuclei and perivitelline sperm. The numbers of perivitelline sperm were counted and Z-stack images were captured under a Nikon Eclipse Ti microscope equipped with a Nikon C2 confocal module (Nikon, Tokyo, Japan).

### *In vitro* fertilization (IVF)

Cauda epididymal spermatozoa extracted from sexually mature wild-type and *Tmem81*^-/-^ males were preincubated in the TYH medium for 2 h at 37 °C, 5% CO_2_ to induce capacitation. Cumulus-oocyte complexes (COCs) were extracted from the oviductal ampulla of superovulated wild-type female mice and were then treated with 330 μg/mL hyaluronidase (Wako, Osaka, Japan) to disperse the cumulus cells or 1 mg/mL collagenase (Sigma-Aldrich, St. Louis, MO) to remove both the cumulus cells and the ZP. Cumulus-intact and cumulus-free eggs were incubated with sperm at a concentration of 2 × 10^5^ sperm/mL in 100 μL TYH medium drops, whereas ZP-free eggs were inseminated at a density of 2 × 10^4^ sperm/mL. After 6 h of incubation, fertilization success was determined by the formation of two pronuclei.

### Sperm–oolemma binding and fusion assays

To investigate the oolemma binding ability, cauda epididymal sperm were preincubated in TYH medium at 37 °C, 5% CO_2_ for 2 h. The capacitated sperm were probed with an anti-IZUMO1 monoclonal antibody (#125) and an Alexa Fluor™ 488-conjugated secondary antibody in TYH medium drops. Meanwhile, COCs were extracted from superovulated wild-type females and treated with 1 mg/mL of collagenase to remove both the cumulus cells and the ZP. The ZP-free eggs were then incubated with the antibody-probed spermatozoa at a density of 2 × 10^4^ sperm/mL. After insemination for 30 min, the sperm–egg complexes were fixed on ice in 0.25% glutaraldehyde in FHM for 15 min, gently washed in fresh FHM medium drops, and stained with Hoechst 33342 to visualize the bound sperm. The total number of sperm bound the oolemma and the number of acrosome-reacted sperm bound to the oolemma were counted under a Keyence BZ-X810 microscope (Keyence, Osaka, Japan). Z-stack images were captured under a Nikon Eclipse Ti microscope equipped with a Nikon C2 confocal module (Nikon, Tokyo, Japan).

For analyzing sperm–egg fusion, wild-type ZP-free eggs were stained with Hoechst 33342 for 15 min at a dilution of 1:10,000, washed thoroughly in fresh TYH medium drops, and incubated with capacitated sperm in TYH medium. After 30 min of insemination, the sperm– egg complexes were fixed on ice in 0.25% glutaraldehyde in FHM, washed in fresh FHM medium drops, and examined under a confocal microscope.

### Immunoprecipitation from zebrafish sperm lysates followed by mass-spectrometry

Per replicate, 1.6 - 4.1 x 10^8^ sperm were collected in Hank’s saline. Prior to lysis, it was pelleted by spinning samples at 800 g for 3 minutes at 4 °C and resuspended in about 30 μL Hank’s saline. For the crosslinked samples, the pellet was resuspended in 1% formaldehyde in Hank’s saline solution and incubated for 10 minutes at room temperature. 100 mM Tris-HCl (pH 8) was added and incubated for 10 minutes to quench the crosslinking reaction before pelleting prior to lysis. For sperm lysis, 2 µL lysis buffer (15 mM NaHEPES pH 7.3, 30 mM NaCl, 1 mM MgCl_2_, 1% Brij35, 1 U/µL benzonase (Merck), 1X complete protease inhibitor (EDTA-free, Roche), 0.2% SDS or 0.5% SDS for crosslinked samples) per 1 x 10^6^ sperm was added and incubated on ice for 1 hour. Insoluble debris was pelleted at 21,000 x g and 4 °C for 10 minutes. The supernatant was used for the immunoprecipitation was performed on the supernatant with Anti-FLAG M2 Magnetic Beads (Sigma-Aldrich) overnight at 4 °C and shaking at 1100 rpm. The beads were washed two times with lysis buffer and three times with wash buffer (15 mM HEPES, 30 mM NaCl) before being transferred to a clean tube to remove any traces of detergent. Prior to mass spectrometric analysis, bound proteins were proteolytically eluted using LysK before tryptic/FASP digestion.

### Targeted mass spectrometry via parallel reaction monitoring (PRM)

For the measurement of relative protein abundance, 2.5 - 3.5 x 10^7^ sperm per biological replicate were collected and lysed in SDT buffer (4% SDS, 100 mM DTT, 100 mM Tris-HCl pH 7.5, 1 mM MgCl_2_) before digestion of nucleic acids with benzonase. After tryptic digestion, the spectra of unique peptides of the proteins of interest (Izumo1, Spaca6, Tmem81) were recorded. Additionally, peptides of six normalization proteins considered constant (Aco2, Hspd1, Mdh1a, Ogdh, Rps27a, St13) were detected. Per protein at least three peptides were measured. Sperm of each genotype was measured in biological triplicates. The sum of the peptide areas of each normalization protein was divided by the average sum across samples. The values were averaged across all normalization proteins to obtain a normalization factor for each sample that served as correction for total protein input. The sum of peptide areas of each protein of interest was scaled by the normalization factor for each sample. The relative protein abundance in each sample is determined by dividing by the average value of the wild-type samples.

### Statistics and reproducibility

All values are shown as the mean ± SD of at least three independent experiments. After examining the normal distribution and variance, the indicated statistical tests and plotting were performed in GraphPad Prism.

---

**Suppl. Fig. S1:**
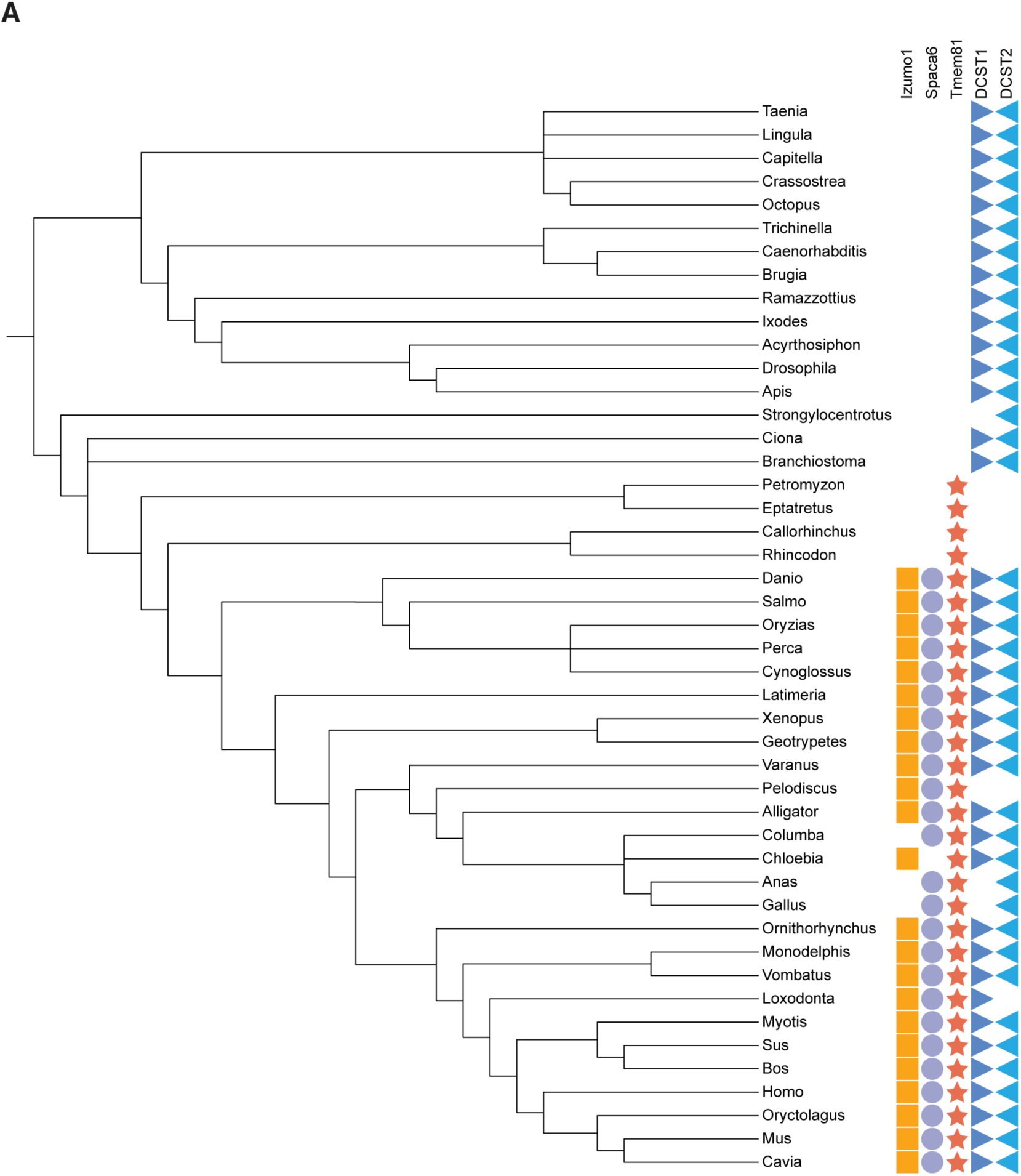
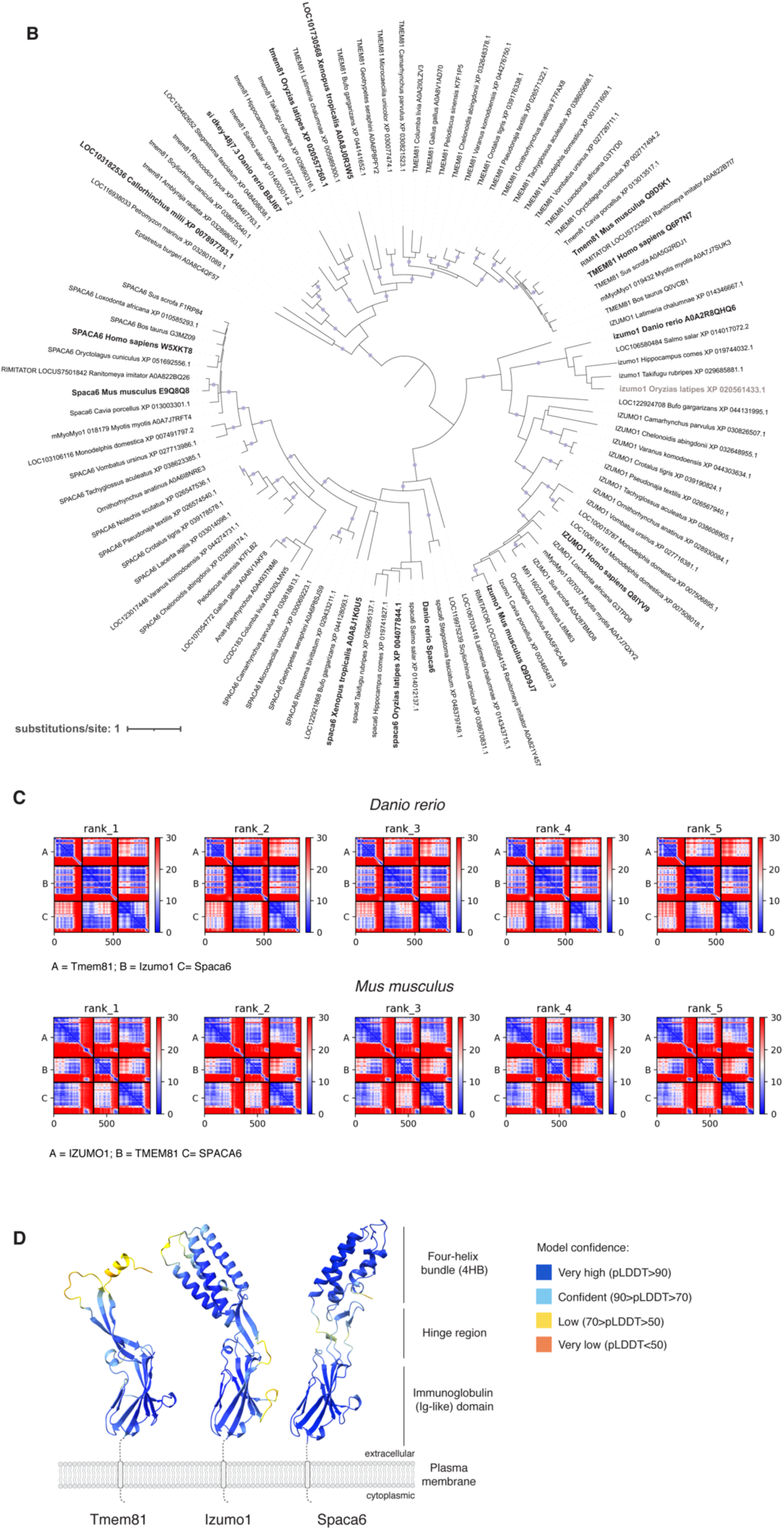
Conservation of vertebrate fertility factors and predicted interaction between Izumo1, Spaca6 and Tmem81. (**A**) Taxonomic tree of Izumo1 (orange square), Spaca6 (blue circle), Tmem81 (red star), Dcst1 (dark blue triangle) and Dcst2 (light blue triangle) across vertebrates and invertebrates. (**B**) Phylogenetic tree of Izumo1, Spaca6, and Tmem81 proteins. Bootstrap values of ≥ 95 are indicated by filled purple circles. Branch lengths represent the inferred number of amino acid substitutions per site, and branch labels are composed of gene name (if available), genus, species, and accession number. (**C**) Predicted Aligned Error (PAE) plots of the top 5 models of the predicted trimeric complex in zebrafish (top) and mouse (bottom). Units: amino acid residues; color bar: Expected position error in Angstroms. (**D**) Tertiary structure predictions of zebrafish Tmem81, Izumo1 and Spaca6. Each amino acid is shaded with per-residue accuracy of the structure (pLDDT) scores. Predictions were generated by AlphaFold2^36^.

**Suppl. Fig. S2:**
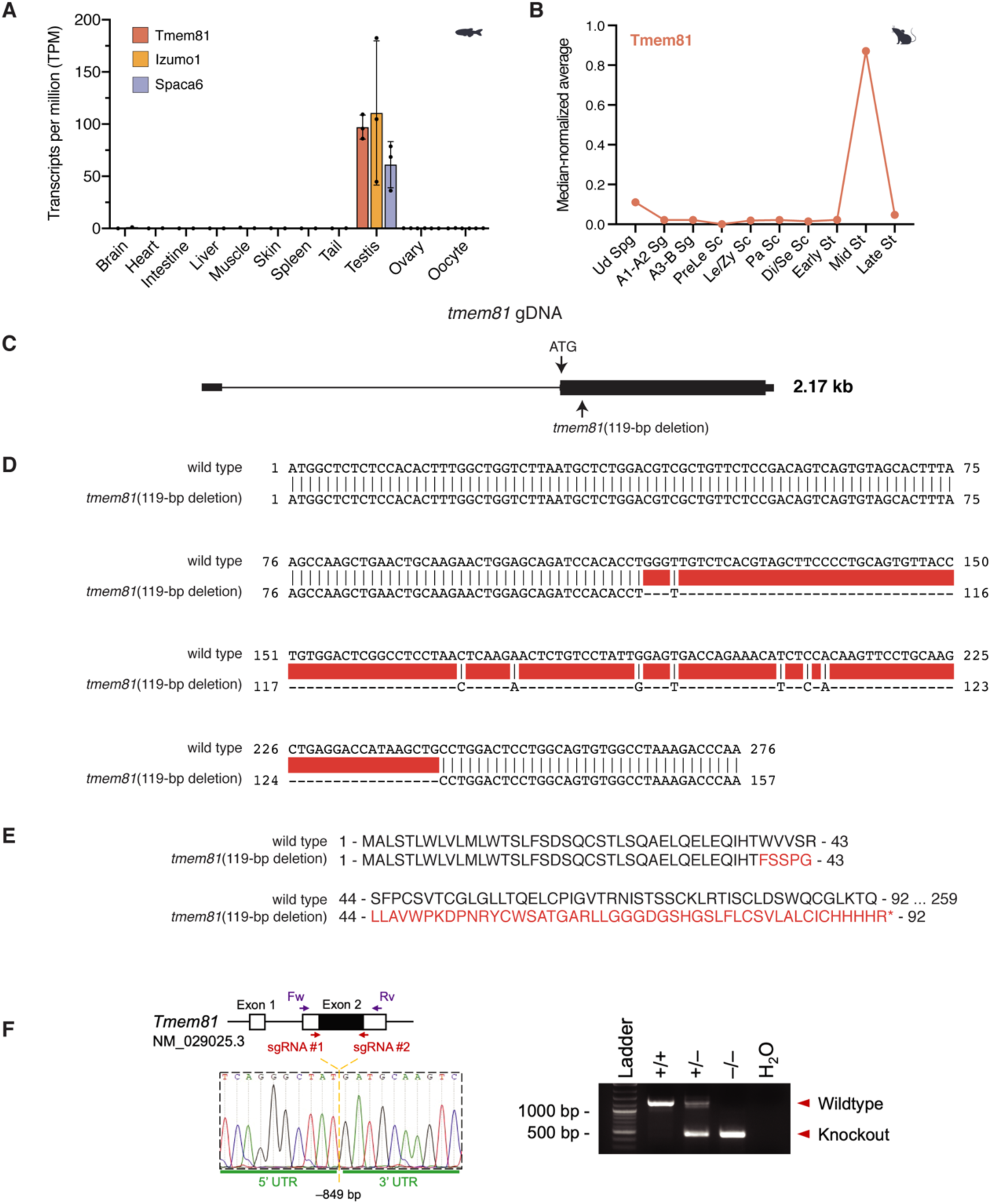
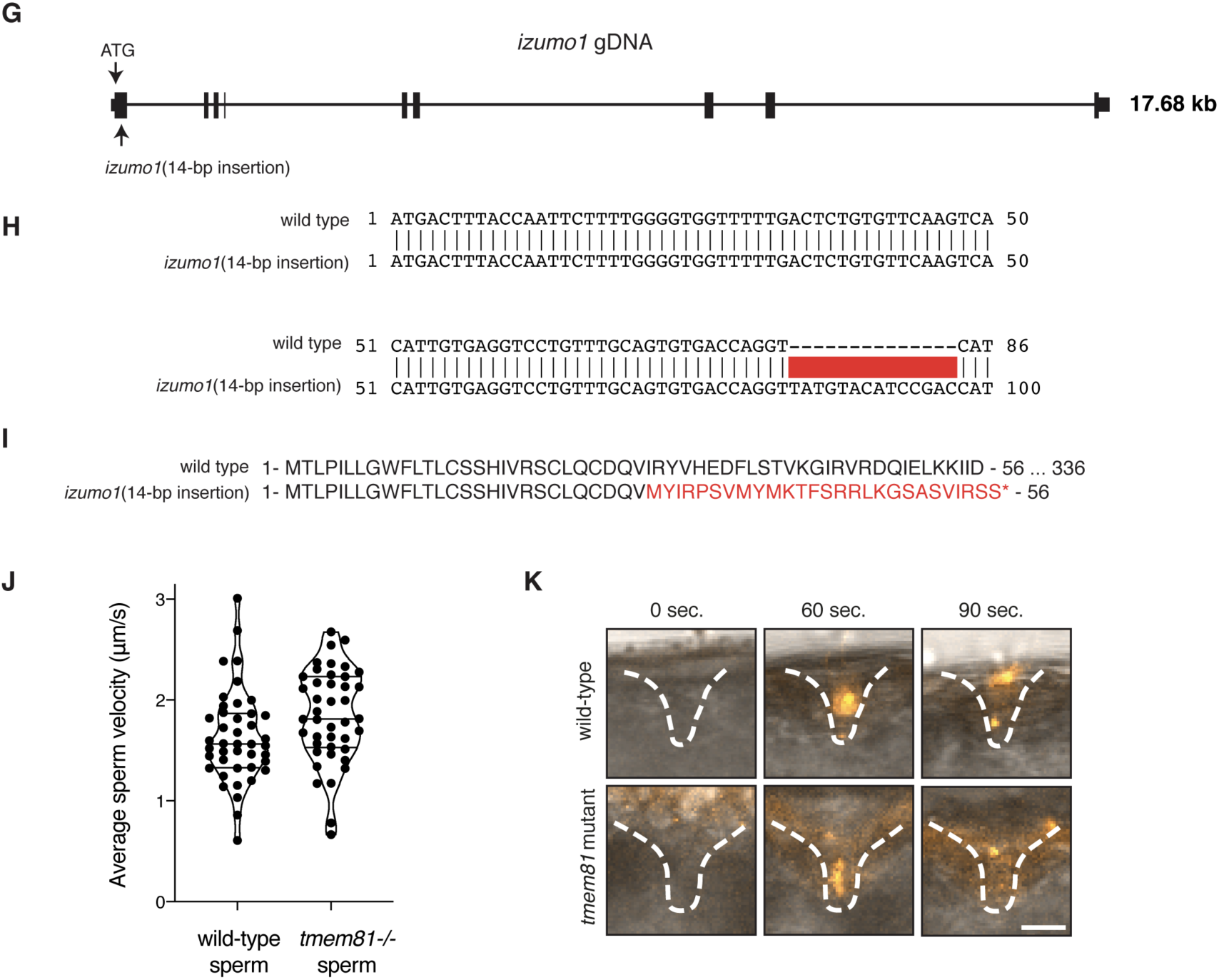
Characterization of expression and function of Tmem81 and Izumo1 in fish. (**A**) Expression of zebrafish *tmem81*, *izumo1* and *spaca6* mRNA in adult tissues. RNA-seq data from^16,27^. (**B**) Median-normalized level of *Tmem81* mRNA expression during mouse spermatogenesis. *Tmem81* is expressed in mid-round spermatids. Ud Spg, undifferentiated spermatogonia; A1-A2 Sg, A1-A2 differentiating spermatogonia; A3-B Sg, A3-A4-In-B differentiating spermatogonia; PreLe Sc, preleptotene spermatocytes; Le/Zy Sc, leptotene/zygotene spermatocytes; Pa Sc, pachytene spermatocytes; Di/Se Sc, diplotene/secondary spermatocytes; Early St, early round spermatids; Mid St, mid round spermatids; Late St, late round spermatids. (**C**) Scheme of the zebrafish *tmem81* locus (ENSDART00000062375.5), indicating the start codon (ATG) and the location of the 119-bp deletion present in the *tmem81* mutant generated in this study. Exons are depicted as boxes, introns as lines. (**D**) Partial cDNA sequence alignment of wild-type and mutant *tmem81*. The *tmem81* mutant has a net deletion of 119-nt highlighted in red. (**E**) Partial amino acid sequence alignment for wild-type (amino acids 1-92) and mutant Tmem81. Deletion in mutant *tmem81* leads to a frameshift that results in a premature stop codon. (**F**) (Left) Knockout strategy of mouse *Tmem81*. *Tmem81* was knocked out using two sgRNAs flanking its coding exon. Forward (Fw) and reverse (Rv) primers targeting the 5’ and 3’ untranslated regions (UTRs), respectively, were used for genomic PCR. A 849-bp deletion was introduced at the locus of *Tmem81*. (Right) Genomic PCR for detecting wild-type and *Tmem81* knockout alleles. (**G**) Scheme of the zebrafish *izumo1* locus (ENSDART00000184278.1), indicating the start codon (ATG) and the location of the 14-bp insertion present in the *izumo1* mutant generated in this study. Exons are depicted as boxes, introns as lines. (**H**) Partial cDNA sequence alignment of wild-type and mutant *izumo1*. The *izumo1* mutant has a 14-nt insertion highlighted in red. (**I**) Partial amino acid sequence alignment for wild-type (amino acids 1-56) and mutant Izumo1. Insertion in mutant *izumo1* leads to a frameshift that results in a premature stop codon. (**J**) Average sperm motility for wild-type and *tmem81* mutant sperm. (**K**) Time-lapse images of sperm approaching the micropyle of wild-type eggs. Representative time series of MitoTracker-labeled wild-type (top) and *tmem81* KO (bottom) sperm (orange) approaching the micropyle (white dashed lines). Scale bar = 50 μm.

**Suppl. Fig. S3:**
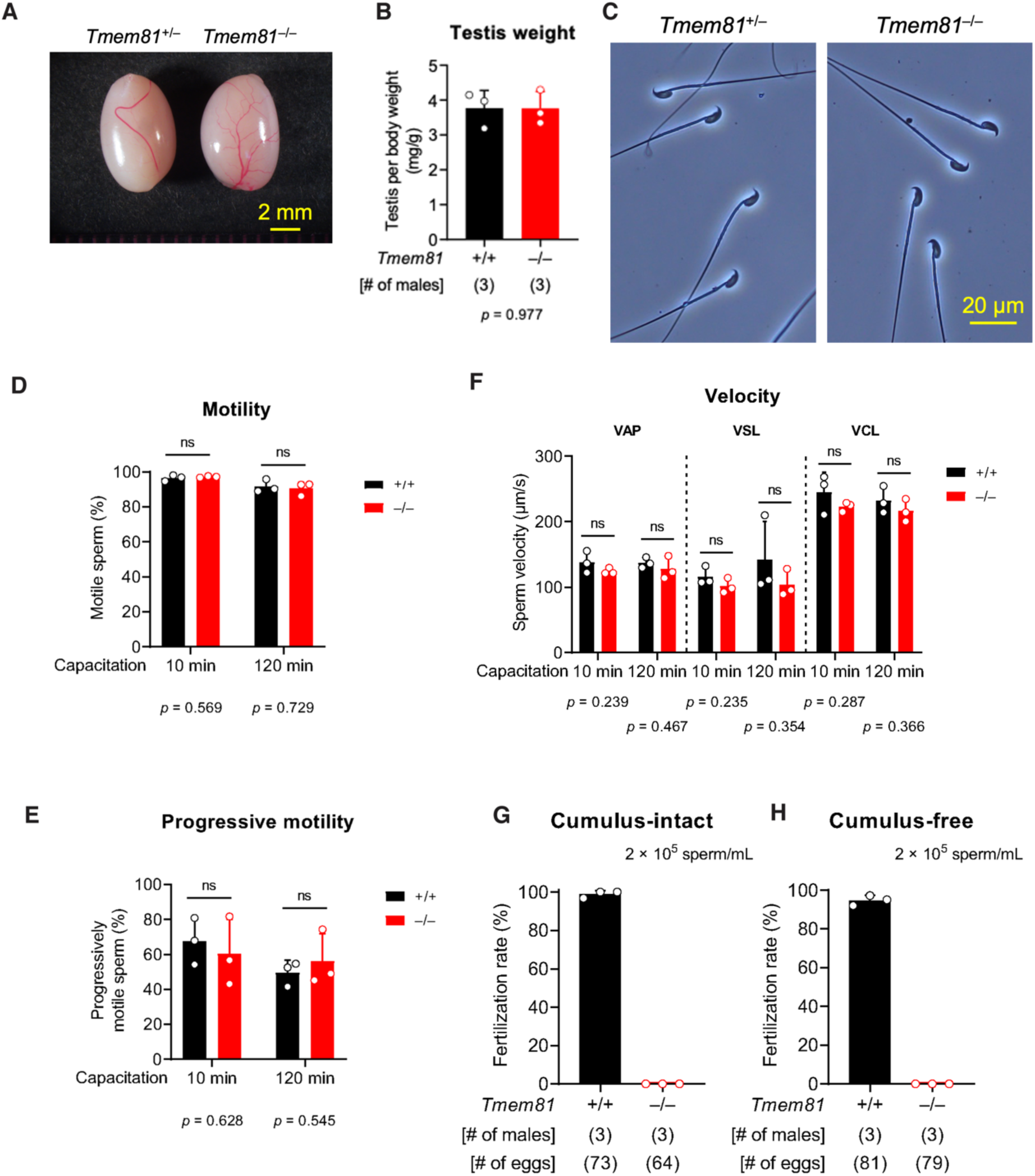
Phenotypic analysis of *Tmem81*^-/-^ male mice. (**A**) Gross appearance of wild-type and *Tmem81*^-/-^ testes. (**B**) Testis weights in wild-type and *Tmem81*^-/-^ males. (**C**) Sperm morphology in wild-type and *Tmem81*^-/-^ males. (**D**) Sperm motility in wild-type and *Tmem81*^-/-^ males. (**E**) Sperm progressive motility in wild-type and *Tmem81*^-/-^ males. (**F**) Sperm swimming velocity in wild-type and *Tmem81*^-/-^ males. (**G**) *In vitro* fertilization between sperm and cumulus-intact eggs. (**H**) *In vitro* fertilization between sperm and cumulus-free eggs.

**Suppl. Fig. S4:**
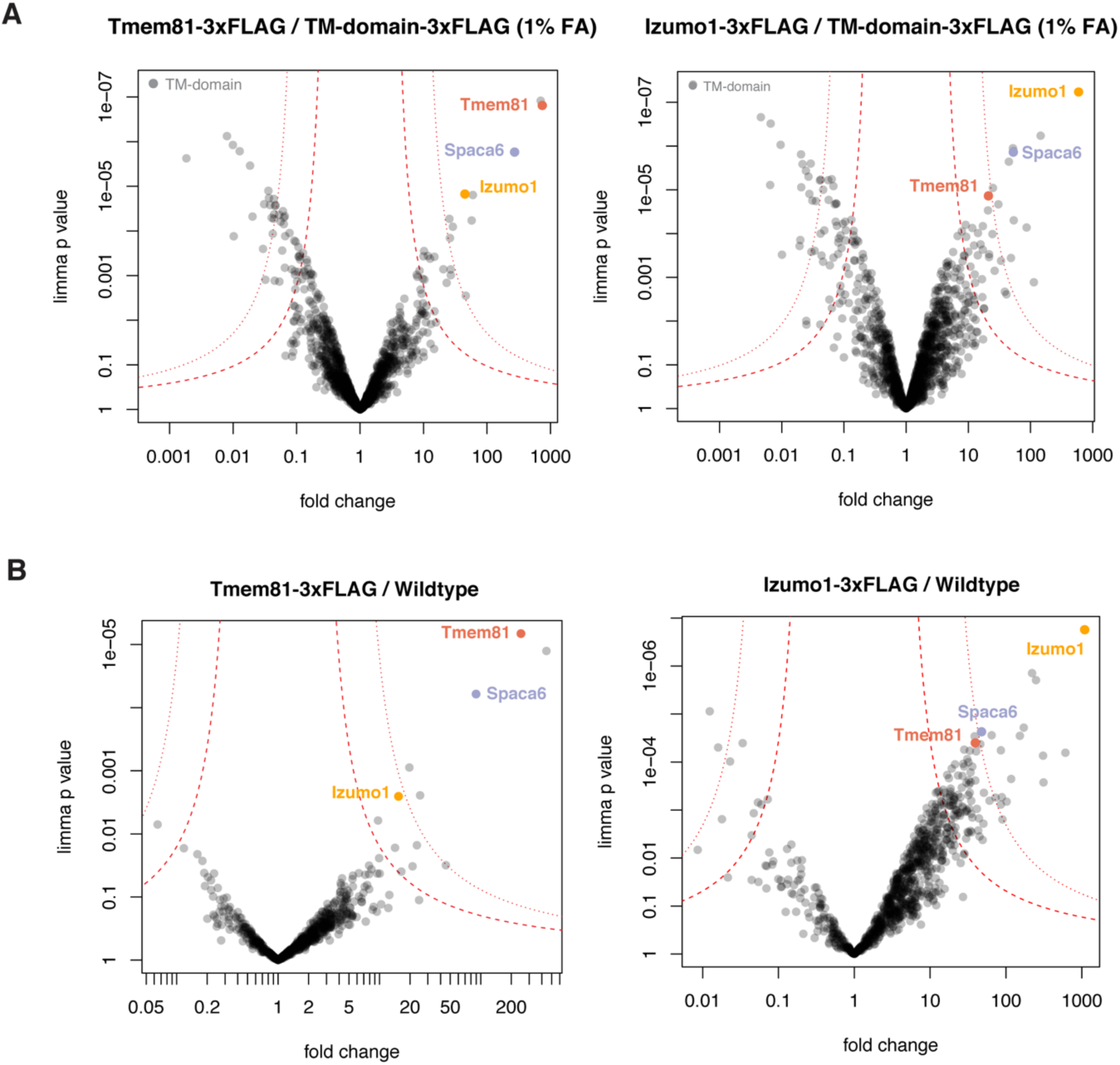
Experimental evidence for the interaction of trimeric complex members Izumo1, Spaca6 and Tmem81 in zebrafish sperm. **(A-B)** Volcano plot of differentially enriched proteins in FLAG co-IPs from Tmem81-3xFLAG-sfGFP **(A)** or Izumo1-3xFLAG-sfGFP **(B)**-expressing sperm versus transmembrane domain (TM)-3xFLAG-sfGFP-expressing sperm. Sperm was lysed after crosslinking with 1% FA (formaldehyde). FLAG co-IPs were performed in triplicates and analyzed by shotgun MS. Trimeric complex members are highlighted. **(C-D)** Volcano plot of differentially enriched proteins in FLAG co-IPs from Tmem81-3xFLAG-sfGFP **(C)** or Izumo1-3xFLAG-sfGFP **(D)** expressing sperm versus wild-type sperm without crosslinking.

**Suppl. Fig. S5:**
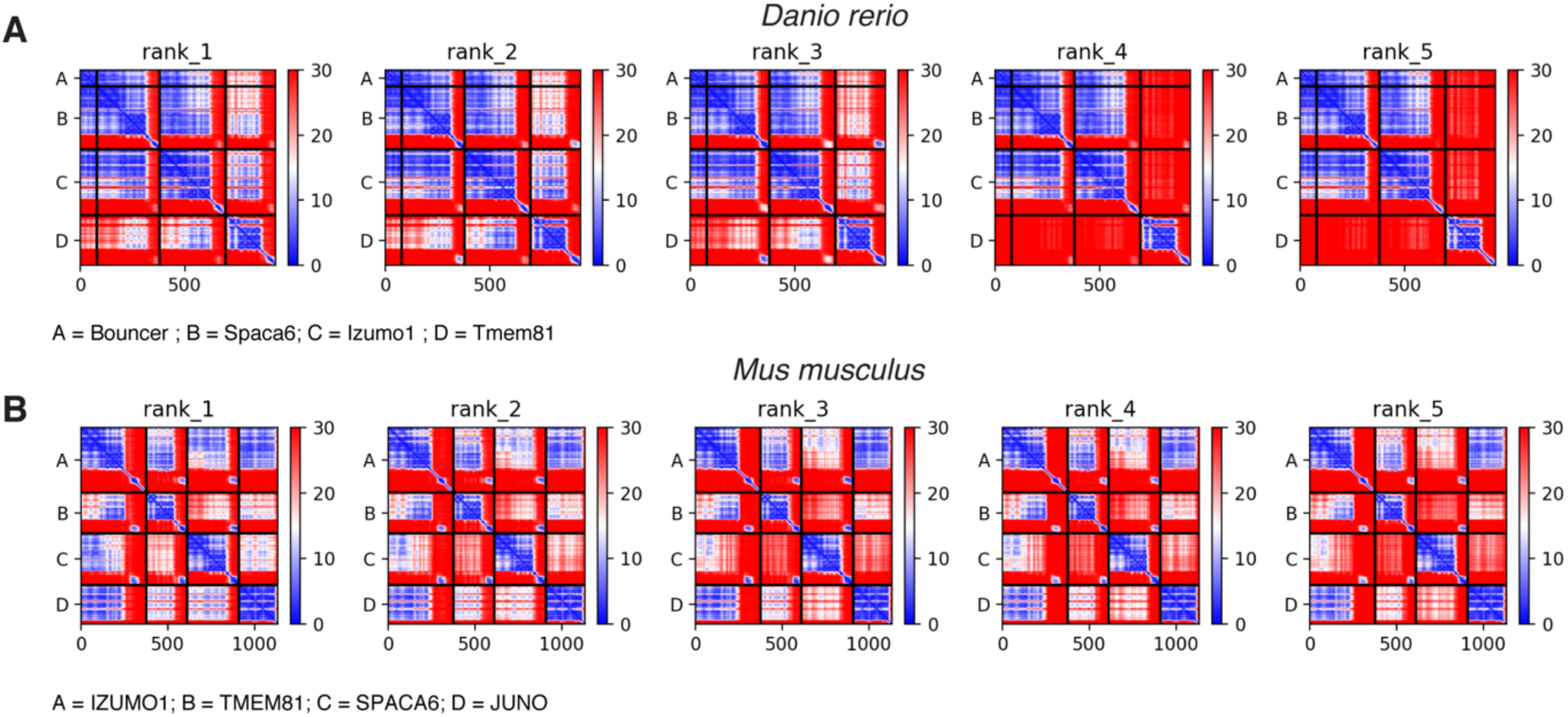
Predicted Aligned Error (PAE) plots of the predicted tetrameric complexes formed between sperm and egg in fish and mice. Predicted Aligned Error (PAE) plots of the top 5 models of the trimeric complex with Bouncer in zebrafish (**A**) and with JUNO in mouse (**B**). Units: amino acid residues; color bar: expected position error in Angstroms.

## REFERENCES

1. Okabe, M. (2018). Sperm-egg interaction and fertilization: Past, present, and future. Biol. Reprod. 99, 134–146. 10.1093/biolre/ioy028.

2. Bianchi, E., and Wright, G.J. (2016). Sperm Meets Egg: The Genetics of Mammalian Fertilization. Annu. Rev. Genet. 50, 93–111. 10.1146/annurev-genet-121415-121834.

3. Deneke, V.E., and Pauli, A. (2021). The Fertilization Enigma: How Sperm and Egg Fuse. Annu. Rev. Cell Dev. Biol. 37, 391–414. 10.1146/annurev-cellbio-120219-021751.

4. Lu, Y., and Ikawa, M. (2022). Eukaryotic fertilization and gamete fusion at a glance. J. Cell Sci. 135, jcs260296. 10.1242/jcs.260296.

5. Inoue, N., Ikawa, M., Isotani, A., and Okabe, M. (2005). The Immunoglobulin Superfamily Protein Izumo is Required for Sperm to Fuse with Eggs. Nature 434, 234–238. 10.1038/nature03362.

6. Satouh, Y., Inoue, N., Ikawa, M., and Okabe, M. (2012). Visualization of the moment of mouse sperm-egg fusion and dynamic localization of IZUMO1. J. Cell Sci. 125, 4985– 4990. 10.1242/jcs.100867.

7. Matsumura, T., Noda, T., Satouh, Y., Morohoshi, A., Yuri, S., Ogawa, M., Lu, Y., Isotani, A., and Ikawa, M. (2022). Sperm IZUMO1 Is Required for Binding Preceding Fusion With Oolemma in Mice and Rats. Front. Cell Dev. Biol. 9.

8. Lorenzetti, D., Poirier, C., Zhao, M., Overbeek, P.A., Harrison, W., and Bishop, C.E. (2014). A transgenic insertion on mouse chromosome 17 inactivates a novel immunoglobulin superfamily gene potentially involved in sperm-egg fusion. Mamm. Genome 25, 141–148. 10.1007/s00335-013-9491-x.

9. Noda, T., Lu, Y., Fujihara, Y., Oura, S., Koyano, T., Kobayashi, S., Matzuk, M.M., and Ikawa, M. (2020). Sperm proteins SOF1, TMEM95, and SPACA6 are required for sperm−oocyte fusion in mice. PNAS, 1–10. 10.1073/pnas.1922650117.

10. Barbaux, S., Ialy-Radio, C., Chalbi, M., Dybal, E., Homps-Legrand, M., Do Cruzeiro, M., Vaiman, D., Wolf, J.P., and Ziyyat, A. (2020). Sperm SPACA6 protein is required for mammalian Sperm-Egg Adhesion/Fusion. Sci. Rep. 10, 1–15. 10.1038/s41598-020-62091-y.

11. Fernandez-Fuertes, B., Laguna-Barraza, R., Fernandez-Gonzalez, R., Gutierrez-Adan, A., Blanco-Fernandez, A., O’Doherty, A.M., Di Fenza, M., Kelly, A.K., Kölle, S., and Lonergan, P. (2017). Subfertility in bulls carrying a nonsense mutation in transmembrane protein 95 is due to failure to interact with the oocyte vestments. Biol. Reprod. 97, 50–60. 10.1093/biolre/iox065.

12. Pausch, H., Kölle, S., Wurmser, C., Schwarzenbacher, H., Emmerling, R., Jansen, S., Trottmann, M., Fuerst, C., Götz, K.U., and Fries, R. (2014). A Nonsense Mutation in TMEM95 Encoding a Nondescript Transmembrane Protein Causes Idiopathic Male Subfertility in Cattle. PLoS Genet. 10. 10.1371/journal.pgen.1004044.

13. Lamas-Toranzo, I., Hamze, J.G., Bianchi, E., Fernández-Fuertes, B., Pérez-Cerezales, S., Laguna-Barraza, R., Fernández-González, R., Lonergan, P., Gutiérrez-Adán, A., Wright, G.J., et al. (2020). TMEM95 is a sperm membrane protein essential for mammalian fertilization. eLife 9, e53913. 10.7554/eLife.53913.

14. Fujihara, Y., Lu, Y., Noda, T., Oji, A., Larasati, T., Kojima-kita, K., Yu, Z., Matzuk, R.M., Matzuk, M.M., and Ikawa, M. (2020). Spermatozoa lacking Fertilization Influencing Membrane Protein (FIMP) fail to fuse with oocytes in mice. PNAS, 1–8. 10.1073/pnas.1917060117.

15. Inoue, N., Hagihara, Y., and Wada, I. (2021). Evolutionarily conserved sperm factors, DCST1 and DCST2, are required for gamete fusion. eLife 49, 1–12.

16. Noda, T., Blaha, A., Fujihara, Y., Gert, K.R., Emori, C., Deneke, V.E., Oura, S., Panser, K., Lu, Y., Berent, S., et al. (2022). Sperm membrane proteins DCST1 and DCST2 are required for sperm-egg interaction in mice and fish. Commun. Biol. 5, 332. 10.1038/s42003-022-03289-w.

17. Kaji, K., Oda, S., Shikano, T., Ohnuki, T., Uematsu, Y., Sakagami, J., Tada, N., Miyazaki, S., and Kudo, A. (2000). The gamete fusion process is defective in eggs of Cd9-deficient mice. Nat. Genet. 24, 279–282. 10.1038/73502.

18. Miyado, K., Yamada, G., Yamada, S., Hasuwa, H., Nakamura, Y., Ryu, F., Suzuki, K., Kosai, K., Inoue, K., Ogura, A., et al. (2000). Requirement of CD9 on the Egg Plasma Membrane for Fertilization. Science 287, 321–324. 10.1126/science.287.5451.321.

19. Le Naour, F., Rubinstein, E., Jasmin, C., Prenant, M., and Boucheix, C. (2000). Severely Reduced Female Fertility in CD9-Deficient Mice. Science 287, 319–321. 10.1126/science.287.5451.319.

20. Rubinstein, E., Ziyyat, A., Prenant, M., Wrobel, E., Wolf, J.P., Levy, S., Le Naour, F., and Boucheix, C. (2006). Reduced fertility of female mice lacking CD81. Dev. Biol. 290, 351–358. 10.1016/j.ydbio.2005.11.031.

21. Bianchi, E., Doe, B., Goulding, D., and Wright, G.J. (2014). Juno is the egg Izumo receptor and is essential for mammalian fertilization. Nature 508, 483–487. 10.1038/nature13203.

22. Aydin, H., Sultana, A., Li, S., Thavalingam, A., and Lee, J.E. (2016). Molecular architecture of the human sperm IZUMO1 and egg JUNO fertilization complex. Nature 534, 562–565. 10.1038/nature18595.

23. Ohto, U., Ishida, H., Krayukhina, E., Uchiyama, S., Inoue, N., and Shimizu, T. (2016). Structure of IZUMO1-JUNO reveals sperm-oocyte recognition during mammalian fertilization. Nature 534, 566–569. 10.1038/nature18596.

24. Kato, K., Satouh, Y., Nishimasu, H., Kurabayashi, A., Morita, J., Fujihara, Y., Oji, A., Ishitani, R., Ikawa, M., and Nureki, O. (2016). Structural and functional insights into IZUMO1 recognition by JUNO in mammalian fertilization. Nat. Commun. 7, 1–9. 10.1038/ncomms12198.

25. Binner, M.I., Kogan, A., Panser, K., Schleiffer, A., Deneke, V.E., and Pauli, A. (2022). The Sperm Protein Spaca6 is Essential for Fertilization in Zebrafish. Front. Cell Dev. Biol. 9, 3695. 10.3389/fcell.2021.806982.

26. Grayson, P. (2015). Izumo1 and Juno: the evolutionary origins and coevolution of essential sperm – egg binding partners. R. Soc. Open Sci. 2, 1–11. 10.1098/rsos.150296.

27. Herberg, S., Gert, K.R., Schleiffer, A., and Pauli, A. (2018). The Ly6/uPAR protein Bouncer is necessary and sufficient for species-specific fertilization. Science 361, 1029–1033. 10.1126/science.aat7113.

28. Gert, K.R.B., Panser, K., Surm, J., Steinmetz, B.S., Schleiffer, A., Jovine, L., Moran, Y., Kondrashov, F., and Pauli, A. (2023). Divergent molecular signatures in fish Bouncer proteins define cross-fertilization boundaries. Nat. Commun. 14, 3506. 10.1038/s41467-023-39317-4.

29. Fujihara, Y., Herberg, S., Blaha, A., Panser, K., and Kobayashi, K. (2021). The conserved fertility factor SPACA4 / Bouncer has divergent modes of action in vertebrate fertilization. PNAS 118, 1–14. 10.1073/pnas.2108777118.

30. Chen, L., Song, J., Zhang, J., Luo, Z., Chen, X., Zhou, C., and Shen, X. (2023). Spermatogenic cell-specific SPACA4 is essential for efficient sperm-zona pellucida binding in vitro. Front. Cell Dev. Biol. 11.

31. Brukman, N.G., Nakajima, K.P., Valansi, C., Flyak, K., Li, X., Higashiyama, T., and Podbilewicz, B. (2022). A novel function for the sperm adhesion protein IZUMO1 in cell– cell fusion. J. Cell Biol. 222, e202207147. 10.1083/jcb.202207147.

32. Inoue, N., Hamada, D., Kamikubo, H., Hirata, K., Kataoka, M., Yamamoto, M., Ikawa, M., Okabe, M., and Hagihara, Y. (2013). Molecular dissection of IZUMO1, a sperm protein essential for sperm-egg fusion. Development 140, 3221–3229. 10.1242/dev.094854.

33. Nishimura, K., Han, L., Bianchi, E., Wright, G.J., de Sanctis, D., and Jovine, L. (2016). The structure of sperm Izumo1 reveals unexpected similarities with Plasmodium invasion proteins. Curr. Biol. 26, R661–R662. 10.1016/j.cub.2016.06.028.

34. Vance, T.D.R., Yip, P., Jiménez, E., Li, S., Gawol, D., Byrnes, J., Usón, I., Ziyyat, A., and Lee, J.E. (2022). SPACA6 ectodomain structure reveals a conserved superfamily of gamete fusion-associated proteins. Commun. Biol. 5, 1–14. 10.1038/s42003-022-03883-y.

35. Tang, S., Lu, Y., Skinner, W.M., Sanyal, M., Lishko, P.V., Ikawa, M., and Kim, P.S. (2022). Human sperm TMEM95 binds eggs and facilitates membrane fusion. Proc. Natl. Acad. Sci. 119, e2207805119. 10.1073/pnas.2207805119.

36. Jumper, J., Evans, R., Pritzel, A., Green, T., Figurnov, M., Ronneberger, O., Tunyasuvunakool, K., Bates, R., Žídek, A., Potapenko, A., et al. (2021). Highly accurate protein structure prediction with AlphaFold. Nature 596, 583–589. 10.1038/s41586-021-03819-2.

37. Evans, R., O’Neill, M., Pritzel, A., Antropova, N., Senior, A., Green, T., Žídek, A., Bates, R., Blackwell, S., Yim, J., et al. (2021). Protein complex prediction with AlphaFold-Multimer (Bioinformatics) 10.1101/2021.10.04.463034.

38. Mirdita, M., Schütze, K., Moriwaki, Y., Heo, L., Ovchinnikov, S., and Steinegger, M. (2022). ColabFold - Making protein folding accessible to all. 2021.08.15.456425. 10.1101/2021.08.15.456425.

39. Evans, R., O’Neill, M., Pritzel, A., Antropova, N., Senior, A., Green, T., Žídek, A., Bates, R., Blackwell, S., Yim, J., et al. (2021). Protein complex prediction with AlphaFold-Multimer. bioRxiv, 2021.10.04.463034. 10.1101/2021.10.04.463034.

40. Peterson, A.C., Russell, J.D., Bailey, D.J., Westphall, M.S., and Coon, J.J. (2012). Parallel Reaction Monitoring for High Resolution and High Mass Accuracy Quantitative, Targeted Proteomics. Mol. Cell. Proteomics MCP 11, 1475–1488. 10.1074/mcp.O112.020131.

41. Sanders, D.W., Jumper, C.C., Ackerman, P.J., Bracha, D., Donlic, A., Kenney, D., Castello-serrano, I., Suzuki, S., Tamura, T., Alexander, H., et al. (2021). SARS-CoV-2 Requires Cholesterol for Viral Entry and Pathological Syncytia Formation. eLife, 1–70.

42. Gert, K.R., Quio, L.E.C., Novatchkova, M., Guo, Y., Cairns, B.R., and Pauli, A. (2021). Reciprocal zebrafish-medaka hybrids reveal maternal control of zygotic genome activation timing. bioRxiv.

43. Shetty, J., Wolkowicz, M.J., Digilio, L.C., Klotz, K.L., Jayes, F.L., Diekman, A.B., Westbrook, V.A., Farris, E.M., Hao, Z., Coonrod, S.A., et al. (2003). SAMP14, a novel, acrosomal membrane-associated, glycosylphosphatidylinositol-anchored member of the Ly-6/urokinase-type plasminogen activator receptor superfamily with a role in sperm-egg interaction. J. Biol. Chem. 278, 30506–30515. 10.1074/jbc.M301713200.

44. Ellerman, D.A., Pei, J., Gupta, S., Snell, W.J., Myles, D., and Primakoff, P. (2009). Izumo is part of a multiprotein family whose members form large complexes on mammalian sperm. Mol. Reprod. Dev. 76, 1188–1199. 10.1002/mrd.21092.

45. Inoue, N., Hagihara, Y., Wright, D., Suzuki, T., and Wada, I. (2015). Oocyte-triggered dimerization of sperm IZUMO1 promotes sperm-egg fusion in mice. Nat. Commun. 6, 1–12. 10.1038/ncomms9858.

46. Loughner, C.L., Bruford, E.A., McAndrews, M.S., Delp, E.E., Swamynathan, S., and Swamynathan, S.K. (2016). Organization, evolution and functions of the human and mouse Ly6/uPAR family genes. Hum. Genomics 10, 10–10. 10.1186/s40246-016-0074-2.

47. Ohno, S. (1970). Evolution by Gene Duplication (Springer) 10.1007/978-3-642-86659-3.

48. Rivera, A.M., and Swanson, W.J. (2022). The Importance of Gene Duplication and Domain Repeat Expansion for the Function and Evolution of Fertilization Proteins. Front. Cell Dev. Biol. 10.

49. Elofsson, A., Han, L., Bianchi, E., Wright, G.J., and Jovine, L. (2023). Deep learning insights into the architecture of the mammalian egg-sperm fusion synapse. bioRxiv. https://doi.org/10.1101/2023.07.27.550195.

50. Krauchunas, A.R., Marcello, M.R., and Singson, A. (2016). The molecular complexity of fertilization: Introducing the concept of a fertilization synapse. Mol. Reprod. Dev. 83, 376–386. 10.1002/mrd.22634.

51. Steinegger, M., and Söding, J. (2017). MMseqs2 enables sensitive protein sequence searching for the analysis of massive data sets. Nat. Biotechnol. 35, 1026–1028. 10.1038/nbt.3988.

52. Altschul, S.F., Madden, T.L., Schäffer, A.A., Zhang, J., Zhang, Z., Miller, W., and Lipman, D.J. (1997). Gapped BLAST and PSI-BLAST: a new generation of protein database search programs. Nucleic Acids Res. 25, 3389–3402. 10.1093/nar/25.17.3389.

53. NCBI Resource Coordinators (2018). Database resources of the National Center for Biotechnology Information. Nucleic Acids Res. 46, D8–D13. 10.1093/nar/gkx1095.

54. The UniProt Consortium (2021). UniProt: the universal protein knowledgebase in 2021. Nucleic Acids Res. 49, D480–D489. 10.1093/nar/gkaa1100.

55. Minh, B.Q., Schmidt, H.A., Chernomor, O., Schrempf, D., Woodhams, M.D., von Haeseler, A., and Lanfear, R. (2020). IQ-TREE 2: New Models and Efficient Methods for Phylogenetic Inference in the Genomic Era. Mol. Biol. Evol. 37, 1530–1534. 10.1093/molbev/msaa015.

56. Katoh, K., and Toh, H. (2008). Recent developments in the MAFFT multiple sequence alignment program. Brief. Bioinform. 9, 286–298. 10.1093/bib/bbn013.

57. Waterhouse, A.M., Procter, J.B., Martin, D.M.A., Clamp, M., and Barton, G.J. (2009). Jalview Version 2—a multiple sequence alignment editor and analysis workbench. Bioinformatics 25, 1189–1191. 10.1093/bioinformatics/btp033.

58. Kalyaanamoorthy, S., Minh, B.Q., Wong, T.K.F., von Haeseler, A., and Jermiin, L.S. (2017). ModelFinder: fast model selection for accurate phylogenetic estimates. Nat. Methods 14, 587–589. 10.1038/nmeth.4285.

59. Hoang, D.T., Chernomor, O., von Haeseler, A., Minh, B.Q., and Vinh, L.S. (2018). UFBoot2: Improving the Ultrafast Bootstrap Approximation. Mol. Biol. Evol. 35, 518–522. 10.1093/molbev/msx281.

60. Letunic, I., and Bork, P. (2021). Interactive Tree Of Life (iTOL) v5: an online tool for phylogenetic tree display and annotation. Nucleic Acids Res. 49, W293–W296. 10.1093/nar/gkab301.

61. Mistry, J., Chuguransky, S., Williams, L., Qureshi, M., Salazar, G.A., Sonnhammer, E.L.L., Tosatto, S.C.E., Paladin, L., Raj, S., Richardson, L.J., et al. (2021). Pfam: The protein families database in 2021. Nucleic Acids Res. 49, D412–D419. 10.1093/nar/gkaa913.

62. Gagnon, J.A., Valen, E., Thyme, S.B., Huang, P., Ahkmetova, L., Pauli, A., Montague, T.G., Zimmerman, S., Richter, C., and Schier, A.F. (2014). Efficient mutagenesis by Cas9 protein-mediated oligonucleotide insertion and large-scale assessment of single-guide RNAs. PLoS ONE 9, 5–12. 10.1371/journal.pone.0098186.

